# Independent noise realizations enable morphologically agnostic image reconstruction in single-photon-sensitive microscopy

**DOI:** 10.64898/2026.09.12.751176

**Authors:** Lisa Cuneo, Alessandro Zunino, Laurent Le, Sofia Agostoni, Saverio Salzo, Luca Calatroni, Massimiliano Pontil, Giuseppe Vicidomini

## Abstract

Advances in single-photon sensitive detectors are rapidly expanding the adoption of photon-counting fluorescence microscopy. Under the Poisson photon-counting statistics, iterative Richardson–Lucy (RL) -type algorithms are statistically optimal for image deconvolution but suffers from a fundamental semi-convergent behaviour: prolonged iterations inevitably amplify noise, requiring heuristic early stopping or regularisation typically based on assumptions about object morphology. Here we present a morphology-agnostic regularisation framework for RL deconvolution that exploits independent noise realisations instead of structural object priors. Preserving the Poisson statistics of the acquisition process, we formulate the Regularized by Noise (RbN): a regularized variational framework and its corresponding iterative minimization algorithm that exploit the statistical consistency of independent noise realisations to distinguish reproducible image features from stochastic noise. The regularisation strength is selected automatically using the Poisson residual whiteness principle, resulting in a fully data-driven reconstruction without heuristic parameter tuning. Photon-timing-resolved systems naturally provide the independent noise realisations exploited by our framework, whereas computational photon splitting provides a statistically equivalent implementation for photon-counting systems. We validate the approach experimentally using photon-timing-resolved confocal microscopy and image scanning microscopy across eight morphologically distinct subcellular targets, and further demonstrate its applicability to photon-counting microscopes through computational photon splitting. Across imaging modalities, detector technologies, biological structures and signal-to-noise regimes, our method eliminates RL semi-convergence, removes sensitivity to the stopping criterion, and consistently outperforms conventional RL while preserving fine structural detail. More broadly, our results establish independent noise realisations as a general source of morphology-agnostic regularisation. Although demonstrated here for SPAD-based laser-scanning microscopy and Poisson statistics, the underlying principle could be extended to other imaging modalities and, more generally, to statistical inverse problems through appropriate noise-specific formulations.

## 1 Introduction

Fluorescence microscopy is an indispensable imaging tool in modern cell biology, providing molecular specificity and sensitivity for investigating the spatiotemporal organization of biomolecules and subcellular structures. A central objective in the development of fluorescence microscopy imaging is to maximize spatial resolution despite the fundamental limitations imposed by optical diffraction and measurement noise. Achieving this goal has relied on the combined development of sophisticated optical designs, advanced photon detection architectures, optimized sample preparation and labeling strategies, and powerful computational reconstruction methods. Together, these advances have transformed fluorescence microscopy, enabling capabilities ranging from super-resolution imaging beyond the diffraction limit [1] to increasingly quantitative and information-rich imaging [2, 3]. In particular, the synergistic development of microscopy hardware and computational reconstruction algorithms has established computational microscopy as a powerful paradigm for extending imaging performance beyond the physical limits of optical hardware alone [4, 5].

Single-photon-sensitive detectors are gaining momentum in fluorescence microscopy [6–8], particularly in laser-scanning microscopy, where conventional photo-multiplier tubes are increasingly being replaced by silicon photomultipliers (SiPMs) [9], hybrid photodetectors (HyDs) [10], point single-photon avalanche diodes (SPADs), and asynchronous read-out SPAD arrays [11, 12]. At the same time, photon-counting quantitative CMOS (qCMOS) cameras [13] and megapixels SPAD arrays [6] are bringing photon-counting imaging to wide-field microscopy. By detecting individual photons with negligible readout noise, these technologies preserve the discrete nature of the photon detection process, enabling photon-counting measurements governed by Poisson statistics. Beyond photon-counting, these detectors differ in the additional single-photon information they preserve. Point SPADs and HyDs provide photon-timing-resolved measurements but limit the sustainable photon flux to the megacount-per-second regime,, whereas SiPMs increase the achievable photon rate to the gigacount-per-second regime at the expense of photon-timing information. Asynchronous read-out SPAD arrays offer a compromise, comprising only a few tens of independently read-out SPAD elements which preserving photon-timing-resolved measurements. Furthermore, they retain the spatial location of every detected photon, enabling detector-array-based super-resolution laser-scanning microscopy [12]. Megapixel SPAD arrays extend this concept to millions of pixels but currently operate mainly in a binary high-speed imaging mode because timestamping every detected photon would require prohibitive data bandwidth. Finally, photon-counting qCMOS cameras provide the highest photon-detection efficiency, although without photon-timing information. Collectively, these detector architectures preserve rich photon-level information, creating new opportunities for computational microscopy.

Within computational microscopy, model-based image reconstruction exploits explicit image formation models and noise statistics to recover sample information implicitly encoded in the measurements. Image deconvolution is one of its most prominent applications: by inverting the blurring introduced by the microscope point-spread function (PSF), it recover spatial frequencies approaching the diffraction limit [14–16]. The Richardson–Lucy (RL) iterative algorithm [17, 18] is the canonical approach for image deconvolution, and has evolved through numerous extensions into a general image reconstruction framework underpinning diverse fluorescence microscopy modalities, including multiview light-sheet microscopy [19] and super-resolution imaging [20, 21]. Derived as an expectation–maximization procedure for maximising the Poisson likelihood of the observed image with respect to the latent object, RL naturally combines the microscope image formation model with photon-counting statistics while preserving the non-negativity of the reconstructed image. Despite its widespread use, RL exhibits a fundamental limitation known as semi-convergence [16, 22]. Initially, the reconstruction sharpens as high-spatial-frequency information is recovered, but continued iterations inevitably amplify high-frequency noise, progressively degrading image quality. In practice, RL must therefore be stopped before this noise-dominated regime is reached. Such early stopping acts as an implicit regularization of the inverse problem; however, the optimal stopping iteration of RL is data-dependent and difficult to determine, limiting reconstruction robustness and reproducibility [23]. A recent study established that this behaviour is mathematically inevitable rather than empirical [24]. Specifically, the Cramér–Rao lower bound (CRLB) for the unbiased estimation of high-spatial-frequency object components diverges as the spatial frequency approaches the diffraction cutoff from below. This implies that RL deconvolution is necessarily ill-convergent.

Two broad classes of regularization strategies have been pursued over the last decades to address the semi-convergence of RL deconvolution. The first class relies on model-based regularization, introducing explicit prior assumptions about the object directly into the optimization functional. Such priors promote spatial smoothness or piecewise regularity (*e.g.,* Tikhonov, total-variation, or Hessian-Schatten norm regularization [25–28]), sparsity in suitable transform domains [29, 30], or combinations thereof [31]. Although effective at stabilizing the inversion, these approaches introduce a bias into the reconstruction. This bias can be beneficial when the underlying specimen is well described by the assumed prior, but may distort genuine features and introduce artefacts when the prior does not match the true, generally unknown, specimen structure. Rather than hand-crafting this structural assumption, a second class of methods learns it from data. This includes plug-and-play schemes, which replace the proximal operator of an explicit regularizer with a pretrained denoiser [32, 33]; algorithm-unrolling approaches, which embed iterative model-based updates within trainable network architectures [34, 35], and end-to-end networks, which directly map degraded measurements to restored images without an explicit forward model or iterative optimization [36]. Although highly successful in fluorescence microscopy, these methods inherit the structural and noise statistics of their training data, and their performance can degrade when specimen morphology or imaging conditions depart from those encountered during training.

Fluorescence microscopy images span morphologies ranging from point-like single molecules to continuous membrane networks, making a universal, morphology-agnostic regularization strategy particularly attractive. Achieving this objective requires a source of regularization that does not depend on prior assumptions about the specimen itself. The emergence of single-photon detector technologies offers an opportunity to revisit this long-standing limitation of Poisson maximum-likelihood reconstruction. Beyond improving measurement sensitivity and faithfully preserving photon-counting statistics, these technologies provide access to richer statistical information that has remained largely unexplored by image reconstruction algorithms. This raises a fundamental question: can the additional statistical information provided by single-photon detectors itself become the source of regularization? More specifically, can the instability of Poisson maximum-likelihood reconstruction be overcome without introducing explicit or implicit assumptions about specimen morphology, instead exploiting only the statistical information intrinsically available within the acquired measurement?

Here, we answer this question affirmatively. A distinctive feature of photon-counting detection is that it enables access to multiple statistically independent realizations of the same underlying object. Provided specimen dynamics are negligible during image acquisition, photon-timing-resolved laser-scanning microscopy, as enabled by detectors such as hybrid photodetectors, point SPADs, and asynchronous read-out SPAD arrays, naturally provides independent realizations by partitioning the detected photon stream at each scan position into temporal windows. Similarly, in large-format SPAD arrays, independent realizations can be obtained by grouping high-speed image frames. In purely photon-counting measurements, such as those acquired with qCMOS cameras or SiPMs, equivalent realizations can instead be generated computationally through stochastic photon splitting [37]. Although each realization contains only a subset of the detected photons and is therefore noisier than the complete image, they all represent independent observations of the same specimen and consequently share the same underlying structure while differing only in their photon-counting noise. We exploit this property by introducing Regularized by Noise (RbN), a variational framework that regularizes Poisson maximum-likelihood reconstruction through statistical consistency among independent realizations of the same measurement. The framework assigns an independent reconstruction variable to each realization and penalizes their variability, allowing regularization to emerge directly from the acquired data rather than from assumptions about specimen morphology. Similar idea has recently emerged in other contexts. In self-supervised image restoration and inverse problems, methods such as Noise2Noise [38], Noise2Self [39], and many related approaches exploit independent or statistically decoupled noisy measurements to train denoising or reconstruction networks without requiring clean targets; we refer the reader to the recent review by Tachella and Davies [40]. More recently, optimization methods based on gradient consensus have shown that agreement between gradients computed from independent measurements can improve convergence and reduce overfitting to realization-specific noise during iterative reconstruction [41]. RbN is conceptually distinct. Rather than using independent observations for network training or to guide the optimization process, it incorporates their statistical agreement directly into the variational formulation of the inverse problem, transforming measurement redundancy into an explicit regularization principle. The regularizer penalizes discrepancy between statistically independent reconstructions while suppressing realization-specific noise. Consequently, RbN requires neither training data nor handcrafted image priors, remaining fully consistent with the Poisson image formation model.

We solve the resulting RbN variational problem using an efficient scaled gradient projection algorithm, belonging to the broader class of variable-metric first-order methods [42–45]. As in any explicit regularization framework, the reconstruction is governed by a parameter that balances data fidelity against regularization. Rather than selecting this parameter empirically, we determine it automatically using the Poisson residual whiteness principle (PRWP) [46–48], which identifies the regularization strength yielding residuals that are statistically consistent with the expected Poisson noise, resulting in a fully data-driven reconstruction framework. We validate the proposed RbN framework on experimental datasets acquired with a custom laser-scanning microscope equipped with an asynchronous read-out SPAD array detector and a dedicated photon-timing-resolved data acquisition module, enabling both direct access to statistically independent realizations. This platform enables validation of RbN for both the classical Poisson image deconvolution problem in conventional confocal microscopy and image reconstruction in image scanning microscopy (ISM) [49–51], a detector-array extension of confocal microscopy that achieves super-resolution [21, 52] and enhanced optical sectioning [53]. We further validate RbN on commercial photon-counting laser-scanning microscopes, including an ISM system based on a photon-counting SPAD array and a confocal system based on SiPM detection. In these systems, statistically equivalent realizations are generated computationally through stochastic photon splitting.

## 2 Results

### 2.1 The regularization by noise framework

The regularization by noise (RbN) framework exploits multiple statistically independent realizations of the same underlying object to regularize inverse imaging problems without relying on assumptions about specimen morphology (Figure 1).

**Figure 1:**
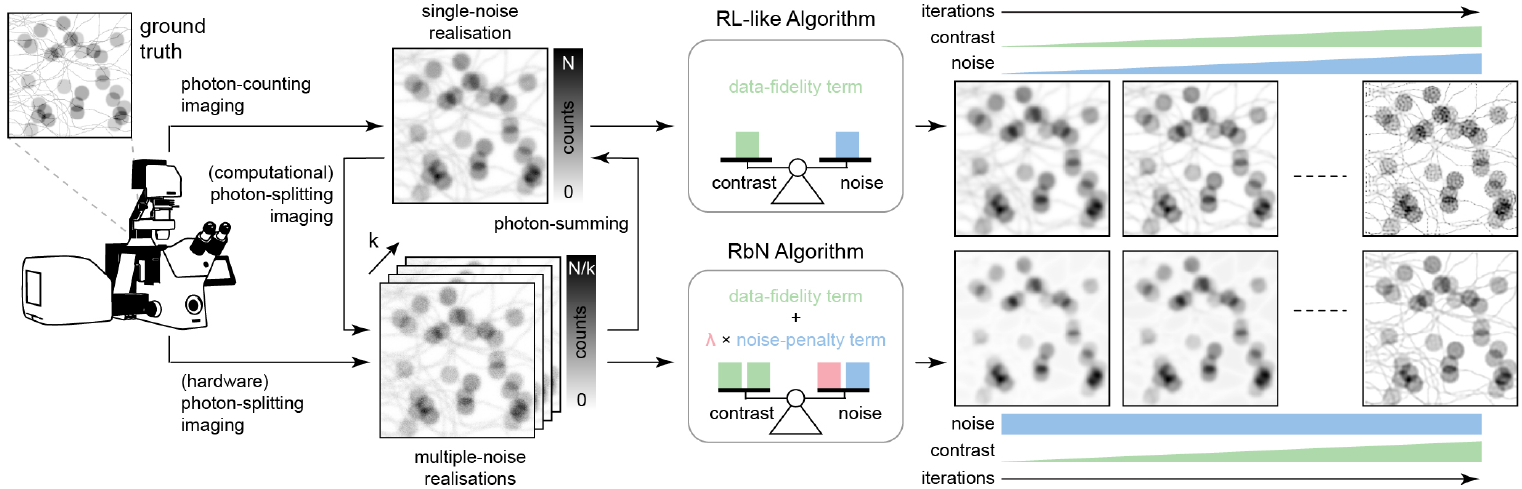
A conventional photon-counting imaging system typically acquires a single Poisson noise realisation of an unknown sample. Contrast-enhancing deconvolution is then commonly performed using Richardson–Lucy (RL)-like iterative algorithms that minimise a Poisson data-fidelity term through an MLEM scheme. Because these algorithms exhibit semi-convergent behaviour, image contrast initially improves but noise is progressively amplified, requiring early stopping based on heuristic criteria. In contrast, the proposed RbN framework exploits multiple independent noise realisations, obtained either experimentally through photon-splitting imaging or synthetically through computational photon splitting. Reconstructions from the different realisations are coupled through a noise-penalty term that promotes proximity to their empirical mean, thereby reducing the empirical variance across reconstructions while preserving the Poisson data-fidelity. As a result, RbN prevents the progressive noise amplification typically observed with RL, enabling progressive contrast enhancement and improved image quality without relying on early stopping. The weighting parameter balancing the data-fidelity and noise-penalty terms is estimated automatically by enforcing the optimal whiteness of the standardised noise residual.

The central idea behind RbN is that independent observations of the same specimen share the same underlying biological structure while differing only in their Poisson noise realization. This statistical redundancy provides a principled way to distinguish signal from noise directly from the experimental data. Features that are consistently recovered across independent realizations are likely to originate from the specimen, whereas discrepancies predominantly reflect stochastic photon-counting fluctuations. RbN incorporates this principle within a variational formulation that combines a realization-specific data-fidelity term with a consistency-based regularizer. The resulting regularization is therefore intrinsically morphology agnostic, depending solely on the statistical consistency of the measurements rather than on assumptions regarding object geometry, smoothness or sparsity. This makes the framework naturally suited to fluorescence microscopy, where biological specimens exhibit remarkably diverse spatial organizations, including isolated puncta, filamentous cytoskeletal networks, membrane systems with piecewise-smooth organization, and densely packed intracellular structures. The only prerequisite of the framework is the availability of statistically independent realizations of the same photon-counting measurement. Such realizations arise naturally in SPAD-based photon-timing-resolved imaging or high-frame-rate binary imaging, and can also be generated computationally through stochastic photon splitting of a single photon-counting image (Figure 1), extending the applicability of RbN beyond dedicated hardware implementations to virtually any photon-counting microscopy modality. By exploiting this statistical redundancy directly within the reconstruction process, RbN replaces the implicit regularization traditionally achieved through early stopping with an explicit regularization. Consequently, reconstruction no longer relies on selecting a heuristic stopping iteration but instead proceeds until convergence of the regularized optimization problem. We formalized this principle by assigning an independent reconstruction variable *u_k_*to each noise realisation *v_k_*, while jointly estimating all reconstructions through a variational formulation that combines a realization-specific Poisson data-fidelity term with a regularization term enforcing statistical consistency among the *K* estimates. The data-fidelity term allows each reconstruction to explain its corresponding measurement according to the physical image formation model, whereas the regularization term penalizes the empirical variance across the realization-specific reconstructions, suppressing features that are not reproducible across independent observations and therefore arising from noise. This leads to the following optimization problem:

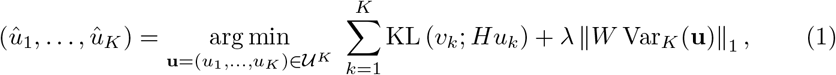

where we seek for an array of solutions (*û*_1_*,…, û_K_*) ∈ *U^K^*, with 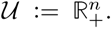 Here 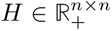 denotes the microscope forward operator, *e.g.,* the convolution matrix encoding the PSF. The regularization strength is controlled by the parameter *ω >* 0, whereas the diagonal weighting matrix 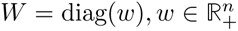 adapts its contribution according to the local reliability of the measurements, allowing spatially varying regularization across the image. The final reconstruction is obtained by combining the realization-specific estimates 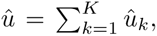 thereby recovering the full photon budget. The first term in eq. (1) preserves the conventional Poisson maximum-likelihood formulation by independently enforcing consistency between each realization *v_k_*and its corresponding forward projected reconstruction *Hu_k_*. On the other hand, the RbN regularization term couples the *K* realisation-specific estimates by penalizing the pixel-wise dispersion through Var*_K_*(**u**), thereby promoting statistical consistency across independent reconstructions. As a result, features consistently recovered across realizations incur little penalty, whereas realization-specific fluctuations are progressively suppressed. We minimized the strongly convex funcitonal in eq. (1) using a positivity-preserving scaled gradient projection algorithm (Supplementary section S1.7) that alternates realization-specific updates with recomputation of their empirical mean entering the variance penalty (algorithm S1). The photon-flux conservation is enforced at every iteration. Unlike unregularized RL, the optimization can be carried to convergence without requiring heuristic early stopping. Because the regularization acts exclusively on statistical consistency across independent realizations rather than on the spatial characteristics of the reconstructed object, it introduces no bias regarding specimen morphology and therefore remains intrinsically morphology agnostic.

### 2.2 Stable, robust, and morphology-agnostic confocal deconvolution

We first validated the RbN framework for image deconvolution in confocal laser-scanning fluorescence microscopy using synthetic datasets with known ground truth (Supplementary Fig. S1), enabling a controlled comparison between RL and RL-RbN. We generated three synthetic phantoms spanning representative biological morphologies, including filamentous cytoskeletal networks, piecewise-smooth membrane-like structures, and a hybrid organization. Across all datasets, conventional RL exhibited the expected semi-convergent behaviour, with reconstruction error initially decreasing before increasing as noise amplification progressively dominated the reconstruction (Supplementary Fig. S1 (a)). The onset of divergence depended strongly on specimen morphology: membrane-like structures entered the unstable regime substantially earlier than sparse filamentous structures, whereas the mixed phantom exhibited intermediate behaviour (Supplementary Fig. S1 (a)). This morphology dependence is consistent with the well known and reported [54] tendency of RL to favour spatially sparse solutions, since filamentous targets are better matched to this implicit sparse bias than extended membrane-like structures. Consequently, early stopping represents a morphology-dependent compromise between structural recovery and noise amplification. Furthermore, selecting the stopping iteration retrospectively using the ground truth to optimize conventional full-reference distortion metrics, including Kull-back–Leibler divergence, PSNR, RMSE, SSIM and its version optimized for microscopy data [55], consistently yielded reconstructions with visible artefacts (Supplementary Figs. S2 to S4). These observations are consistent with the perception–distortion tradeoff, whereby minimizing image distortion does not necessarily maximize perceptual reconstruction quality, irrespective of the distortion metric employed [56]. We next applied RL-RbN using a fixed regularization parameter selected heuristically to suppress noise amplification. Unlike conventional RL, RL-RbN converged smoothly towards a stable solution for all three phantoms without exhibiting semi-convergent behaviour, yielding reconstructions that more faithfully reproduced the underlying structures while remaining free from noise overfitting (Supplementary Fig. S1 (a)). Although retrospectively early-stopped RL consistently achieved lower distortion than RL-RbN according to all full-reference metrics, this advantage relied on selecting the stopping iteration using the ground truth, an oracle condition that is unavailable in real experimental settings. Visual inspection nevertheless revealed inferior structural recovery and the presence of noticeable reconstruction artefacts (Supplementary Fig. S1 (a), Supplementary Figs. S2 to S4), consistent with the perception–distortion trade-off [56] discussed above. To complement conventional distortion metrics, we therefore evaluated reconstruction robustness using a ground-truth-free criterion, quantified from the variability of repeated reconstructions obtained from independent acquisitions of the same specimen [24]. According to this robustness criterion, RL-RbN consistently exhibited substantially lower reconstruction variability than conventional RL across all three synthetic morphologies (Supplementary Fig. S1 (b)), confirming that the proposed regularization effectively suppresses noise sensitivity independently of specimen morphology.

We next validated the RbN framework on experimental photon-counting confocal microscopy using a custom laser-scanning microscope equipped with a SPAD-array detector and dedicated photon-timing-resolved acquisition electronics. By partitioning the photons detected during each pixel dwell time into multiple independent temporal windows, the acquisition directly generates the statistically independent noise realizations required by RbN. The number of realizations, *K*, controls the balance between statistical redundancy and the photon budget available to each realization. Increasing *K* progressively improved the quality of the RbN reconstruction by strengthening the statistical consistency enforced by the regularization. However, excessive photon partitioning, in addition to increasing the computational cost, reduced the photon budget of the individual realizations, causing the regularization to dominate the data-fidelity term and progressively limiting the degree of deconvolution (Supplementary Fig. S5). Unless otherwise stated, all subsequent experiments were therefore performed using *K*=8, which provided an effective compromise between reconstruction quality and computational complexity. Because RbN is an explicit regularization framework, its performance depends on the choice of the regularization parameter *ω*, which balances deconvolution against noise suppression. Rather than selecting this parameter empirically, as in the synthetic validation, we estimated it directly from the acquired data using the Poisson residual whiteness principle (PRWP) [46–48]. PRWP evaluates the standardized residual between the acquired image and the corresponding forward projection of the reconstructed image (Figure 2 (a)). The rationale is that a correctly regularized reconstruction should explain all reproducible image structures, leaving only spatially uncorrelated Poisson fluctuations in the residual. PRWP therefore identifies the optimal regularization strength as the value of *ω* that minimizes the spatial autocorrelation of the standardized residual, providing a fully data-driven criterion for parameter selection. We evaluated this strategy using confocal images of fluorescently labelled F-actin acquired at three signal-to-noise ratios (Figure 2 (b)). For every acquisition condition, the whiteness metric exhibited a well-defined minimum. Consistent with physical expectations, the selected *ω* increased monotonically as the photon budget decreased, indicating that noisier measurements require stronger regularization (Figure 2 (c)). Underestimating *ω* produced RL-like reconstructions at late iterations, with enhanced contrast accompanied by pronounced noise overfitting. Conversely, overestimating *ω* suppressed the deconvolution process, yielding oversmoothed images that largely retained the blurring appearance of the raw data (Figure 2 (d)). Across all imaging conditions, the PRWP-selected regularization parameter consistently achieved a compromise between structural recovery and noise suppression (Supplementary Fig. S6).

**Figure 2:**
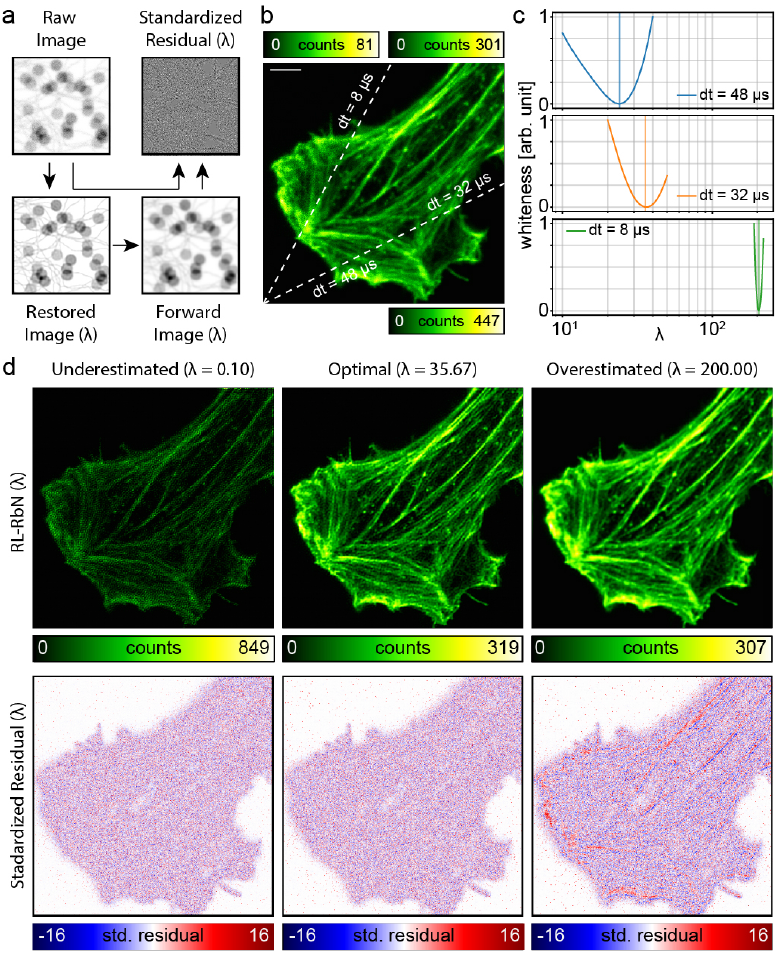
Automatic selection of the RbN regularization parameter through the Poisson Residual Whiteness Principle. (a) Computation of the standardised residual for a given regularisation parameter *ω*. Each RbN reconstruction is mapped back to the measurement space through the forward imaging model *v* = *Hu*, and the corresponding *ω*-dependent standardised residual is computed from the raw image. (b) Representative blurred and noisy CLSM image for the medium-SNR condition (d*t* =32 µs) of fluorescently labelled F-actin (phalloidin–Alexa Fluor 647) acquired using a SPAD-array detector in photon-counting mode with *N*_bin_ = 64 temporal bins per pixel. (c) The residual whiteness metric as a function of *ω* for three acquisition conditions (d*t* = 48, 32, and 8 µs), illustrating that the optimal regularisation weight increases as the photon budget decreases. Vertical lines indicate the automatically selected value of *ω*. (d) Middle row: RbN reconstructions of the medium-SNR dataset obtained with an underestimated (left), optimal (centre), and overestimated (right) value of *ω*. Underestimating *ω* yields RL-like behaviour, with enhanced contrast accompanied by progressive noise overfitting, whereas overestimating *ω* produces oversmoothed reconstructions. The PRWP-selected value provides the best compromise between contrast enhancement and noise suppression. Bottom row: Corresponding standardised residuals. Excessive regularization leaves structured information in the residual, indicating that image content has not been fully explained by the reconstruction. The residual associated with the optimal parameter is spatially white. Although the residuals obtained with underestimated and optimal *ω* appear visually similar after forward projection, the autocorrelation metric detects residual correlations and selects the optimal regularization strength. Scale bar: 5 *µ*m.

Having established a fully data-driven strategy for selecting the regularization parameter, we next compared RL-RbN with conventional RL on the same F-actin datasets (Figure 3, Supplementary Fig. S6). Conventional RL exhibited the expected progressive semi-convergent behaviour: image contrast initially improved, but prolonged iteration progressively amplified structured noise, generating speckle-like artefacts that increasingly obscured the actin network. In practice, this behaviour necessitates conservative early stopping, inevitably sacrificing structural recovery to limit noise amplification (Figure 3 (a)). In contrast, RL-RbN converged smoothly towards a stable solution without exhibiting semi-convergence (Figure 3 (a)). Explicit regularization prevents progressive noise overfitting while allowing continued recovery of structural information. Individual actin filaments appeared sharper and more continuous, neighbouring structures were better separated, and the image background remained largely free of reconstruction artefacts (Figure 3 (a)). For a fair comparison, conventional RL was stopped at the iteration automatically selected by the same PRWP criterion (Supplementary Fig. S7), yielding the reconstruction denoted as RL optimal. Although this objectively selected stopping point substantially reduced noise overfitting, RL-RbN consistently achieved higher filament contrast, more complete structural recovery and a cleaner background (Figure 3 (b)). These results demonstrate that explicit RbN regularization enables superior reconstruction even when conventional RL is stopped at its data-driven optimum (Figure 3 (b), Supplemeng-tary Fig. S8). An additional outcome is that the PRWP provides a principled, fully automatic stopping criterion for conventional RL (Supplementary Figs. S7, S8). We next quantified reconstruction robustness using the ground-truth-free metric introduced for synthetic experiments, computed from repeated independent acquisitions of the same specimen. Conventional RL displayed rapidly increasing reconstruction variability with iteration number, producing highly structured variability maps that closely followed the specimen morphology, a hallmark of noise overfitting. By contrast, RL-RbN maintained consistently low variability with little residual structural content throughout the iterative process (Figure 3(a)). This trend was consistent across all signal-to-noise conditions (Figure 3(c)). Even at the lowest photon budget, RL-RbN achieved lower reconstruction variability than conventional RL at the highest photon budget, demonstrating that explicit noise-based regularization can compensate more effectively for photon-limited imaging than simply increasing the number of detected photons (Figure 3(c)). The benefit of RL-RbN over optimally early-stopped RL was also dependent on the signal-to-noise ratio (Supplementary Fig. S6). At high photon budgets, the PRWP-selected stopping iteration for RL occurred relatively late, allowing substantial recovery of structural information before the onset of noise amplification. As the photon budget decreased, however, the optimal stopping point shifted progressively towards earlier iterations, forcing conventional RL to terminate before fully recovering fine structural details. Consequently, the performance gap between RL-RbN and optimally stopped RL increased under photon-limited conditions, highlighting the ability of explicit RbN regularization to maintain effective deconvolution where early stopping becomes increasingly restrictive (Supplementary Fig. S6).

**Figure 3:**
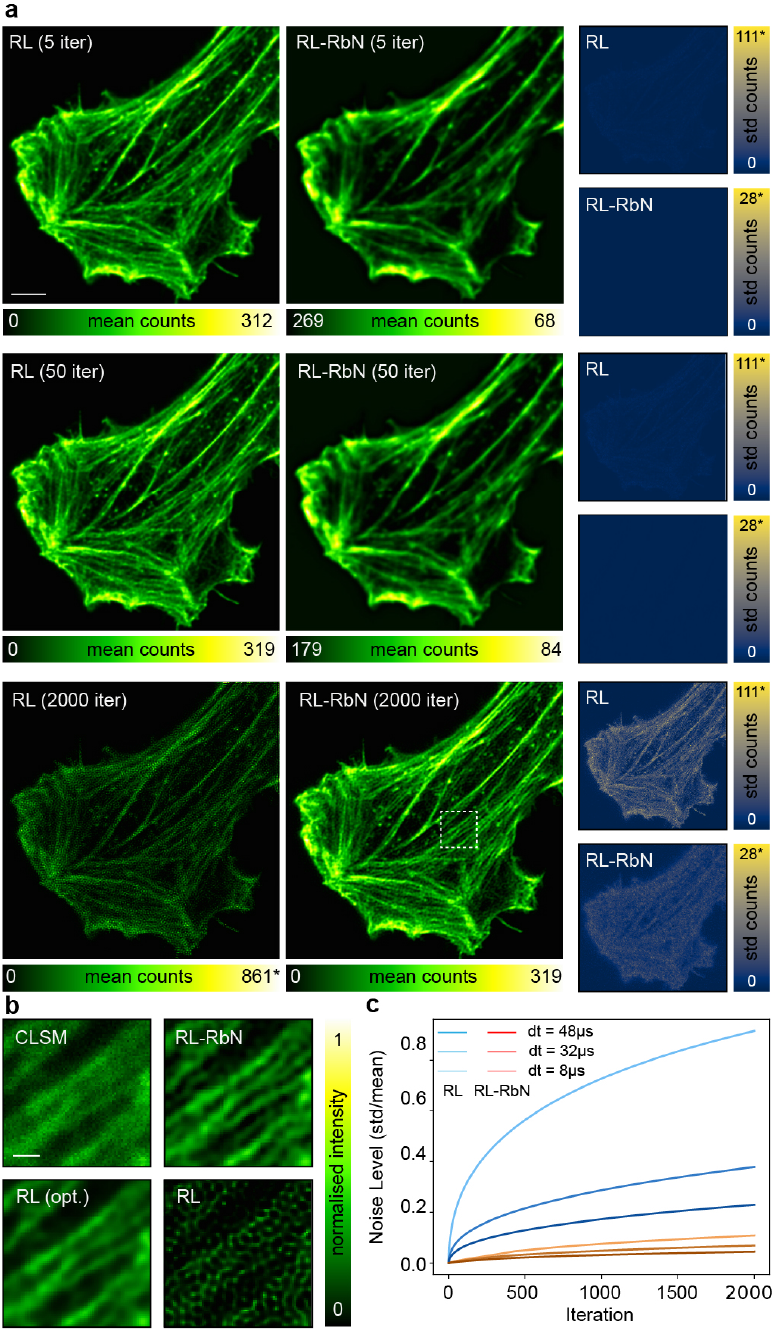
Noise stability of RL and RbN during iterative reconstruction. **(a)** Reconstructions obtained with RL and RL-RbN after 5, 50 and 2000 iterations.. The two left columns show representative reconstructions for the medium-SNR acquisition condition (d*t* = 32 *µ*s), whereas the right column reports, for each procedure, the corresponding standard deviation computed over *L* = 5 repetition of the same experiment with independent noise realizations. In RL, image contrast initially improves but noise is progressively amplified with the number of iterations. In contrast, RL-RbN, with the regularisation parameter automatically selected by the residual whiteness principle (RWP), effectively suppresses noise amplification while preserving contrast enhancement. The asterisk (*) indicates that the displayed maximum corresponds to the 99.9th percentile of the pixel-intensity distribution.**(b)** Magnified regions indicated in (a) comparing the raw confocal image (CLSM), the RL-RbN reconstruction, the PRWP-selected early-stopped RL reconstruction (RL-opt), and the converged RL solution (2000 iterations), highlighting improved structural recovery without noise amplification. **(c)** Evolution of the reconstruction noise level, defined as the ratio between the average standard deviation and the average signal, as a function of iteration number, for three acquisition conditions (d*t* = 48, 32, 8 µs). RL-RbN maintains substantially lower noise growth than conventional RL across all photon budgets. Scale bars: 5 µm (a), 1 µm (b).

To further assess the morphology independence of the proposed regularization, we applied RL-RbN to eight subcellular targets exhibiting markedly different biological organizations: the outer mitochondrial membrane protein TOMM20, forming a fragmented tubular network; the intermediate filament vimentin, arranged as an extended meshwork; the microtubule marker *ε*-tubulin, organized into long, densely packed parallel fibres; the heterochromatin mark histone H3K9me3, distributed in spatially heterogeneous chromatin domains within the nucleus; the nuclear pore complex component NUP153, forming a dense annular array along the nuclear envelope; clathrin-coated pits, appearing as small, isolated diffraction-limited puncta; F-actin, organized into an extended network of intersecting filaments and bundles; and lamin B1, delineating the smooth, continuous nuclear lamina (Figure 4, Supplementary Fig. S10). Across all eight structures, RL-RbN consistently resolves finer structural detail and improves signal-to-noise ratio compared to both the raw CLSM acquisition and standard RL at its optimal stopping iteration, while avoiding the noise amplification observed in conventional RL at late iteration. This versatility is a direct consequence of the agnostic design of the RbN regularization: the variance penalty encodes no information about the spatial structure of the object, acting exclusively on noise-driven inconsistencies across independent noise realizations. The method therefore generalizes across samples and various SNR regime, without introducing morphology-specific reconstruction artefacts, with no modification to any parameter beyond *ω*, which is itself selected automatically by the PRWP, making it broadly applicable to diverse biological specimens and imaging modalities.

**Figure 4:**
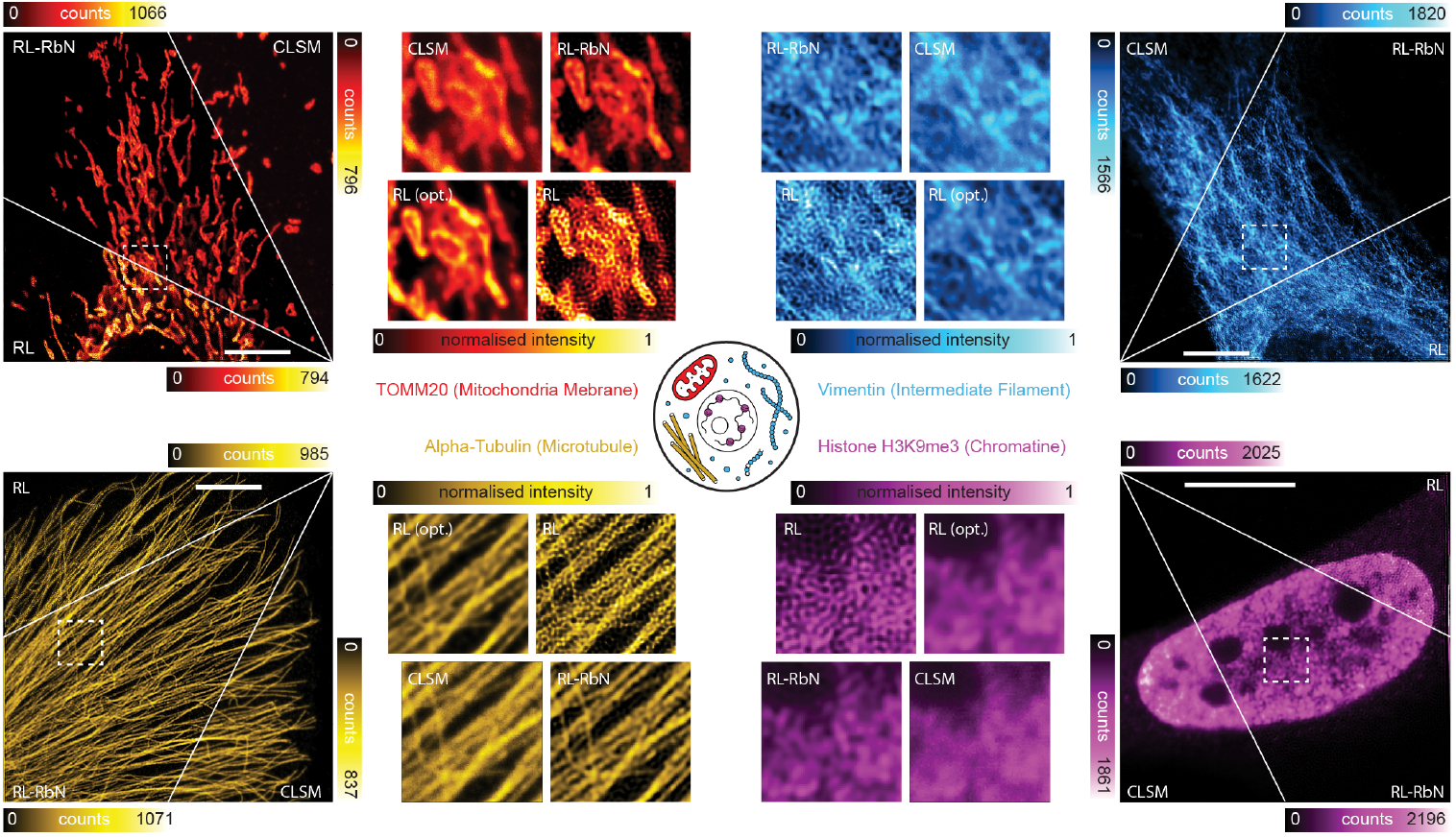
Versatlity of RbN. RL-RbN was applied to CLSM images of four subcellular targets representing morphologically distinct biological organizations: the outer mitochondrial membrane protein TOMM20 (red); the intermediate filament vimentin (cyan); the microtubule marker *ε*-tubulin (yellow); and the heterochromatin mark histone H3K9me3 (magenta). For each target, the full field of view is displayed as a diagonal split comparing the raw CLSM (one triangle) with the RL and RL-RbN reconstructions (other triangle); magnified insets compare CLSM, RL-RbN, RL at its empirically optimal stopping iteration [RL (opt.)], and over-iterated RL. Scale bars: 5 µm (a).

Although the present results were obtained using the independent realizations naturally provided by the photon-timing-resolved acquisition of the SPAD detector, this hardware capability is not required by the RbN framework. Whenever measurements follow Poisson photon-counting statistics, statistically indipendent and identically distribuited realizations can be generated computationally from a single acquisition through stochastic photon splitting, yielding reconstruction performance comparable to the hardware implementation (Supplementary Fig. S9). Consequently, RbN is readily applicable to any photon-counting microscopy platform without dedicated hardware support. To assess the extendability of RbN beyond our custom SPAD-array platform, we applied RL-RbN to photon-counting CLSM datasets acquired with a commercial Evident photon-counting confocal microscope (Supplementary Fig. S14). Since these acquisitions did not provide the intra-pixel temporal partition available on our custom system, the statistically independent realizations required by RbN were generated computationally through stochastic photon splitting. Across *ε*-tubulin and TOMM20-labelled mitochondria, RL-RbN enhanced structural contrast and recovered finer details while suppressing the progressive noise amplification characteristic of prolonged RL iterations.

The same reconstruction framework and automatic PRWP-based parameter selection were used without detector- or specimen-specific modifications, supporting the transferability of RbN across different photon-counting detector architectures.

### 2.3 Regularization by noise extends to ISM reconstruction

Image scanning microscopy (ISM) [49–51] is rapidly emerging as an alternative to conventional confocal microscopy following the introduction of fast detector-array technologies, first through Airyscan [57] and more recently through asynchronous-readout SPAD arrays [52]. By replacing the single confocal detector with a detector array, ISM simultaneously records multiple effective small-pinhole confocal images. Unlike conventional confocal microscopy, where improving resolution requires reducing the pinhole diameter at the expense of photon efficiency, ISM preserves essentially the full detected photon budget while computationally recovering the theoretical twofold lateral resolution enhancement expected for an infinitesimal confocal pinhole. Recent developments have further extended ISM through super-resolution sectioning ISM (s^2^ISM) [53], a maximum-likelihood reconstruction framework that simultaneously achieves twofold lateral resolution enhancement and optical sectioning (see Supplementary section S1.4). We therefore integrated the proposed RbN framework into the s^2^ISM reconstruction algorithm (see Supplementary section S1.5 and Supplementary section S1.6). This represents a substantially more demanding inverse problem than confocal deconvolution, as the forward operator jointly models all detector images while accounting for out-of-focus fluorescence from a three-dimensional specimen to reconstruct a single optical section. Successful integration of RbN within this more complex variational framework demonstrates that the proposed regularization is independent of the underlying image formation operator and therefore generalizes beyond conventional image deconvolution. Compared with conventional confocal microscopy, s^2^ISM-RbN simultaneously improved lateral resolution and optical sectioning while preserving the high photon efficiency of detector-array ISM (Figure 5). Relative to standard s^2^ISM, explicit RbN regularization eliminated progressive noise overfitting, allowing the optimization to recover fine structural information. Individual actin filaments appeared sharper and more continuous, neighbouring structures were better separated, and the out-of-focus background was further suppressed without introducing reconstruction artefacts. Although applying RbN to the closed-pinhole confocal reconstruction also improved image quality, substantially larger gains were achieved when it is combined with s^2^ISM (Figure 5). This behaviour is consistent with the fundamental principle of ISM, which exploits the complementary information collected by the detector array to recover additional spatial information while preserving the full detected photon budget. These improvements were achieved despite the substantially lower photon budget available in each individual detector image, highlighting the robustness of the proposed regularization under highly photon-limited conditions. To assess the generality of the approach, we next applied s^2^ISM-RbN to the same eight subcellular targets analysed in the confocal experiments, namely TOMM20-labelled mitochondria, vimentin, *ε*-tubulin and H3K9me3-labelled chromatin (Figure 6), and the nuclear pore complex component, clathrin-coated pits, F-actin and lamina (Supplementary Fig. S12). Across all structures with different spatial organizations of the specimens, s^2^ISM-RbN consistently preserved the super-resolution and optical-sectioning capabilities of s^2^ISM while suppressing the progressive noise amplification characteristic of RL-like optimization. Direct comparison with the corresponding RL-RbN reconstructions further highlights the additional resolution and sectioning provided by the ISM acquisition while preserving the benefits of RbN regularization (Supplementary Fig. S11). Equivalent performance was obtained using computational photon splitting in place of hardware-generated realizations (Supplementary Fig. S13). Finally, we evaluated s^2^ISM-RbN on datasets acquired with a commercial Nikon microscope equipped with the NSPARC spatial-array detector (Supplementary Fig. S15). The datasets comprised an mouse kidney cryosection and Purkinje cells. For these datasets, the statistically independent realizations required by RbN were generated computationally from the photon-counting detector images. In both samples, s^2^ISM-RbN improved the definition of fine structures while suppressing background and reconstruction noise relative to the confocal image.

**Figure 5:**
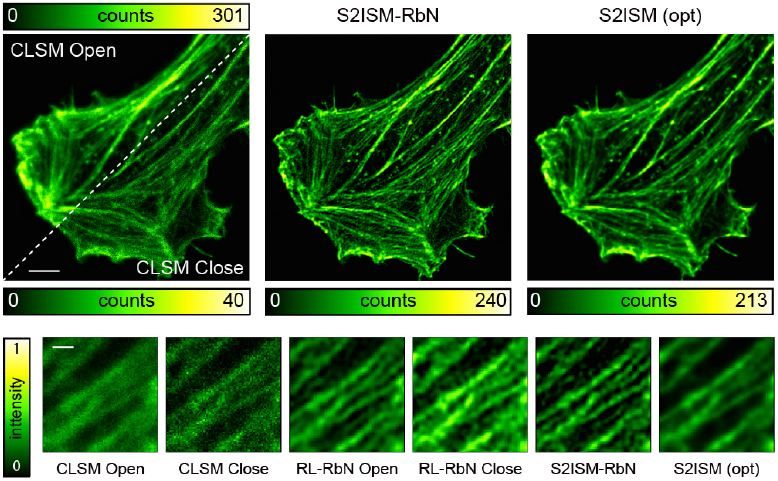
Application of RbN to ISM. Representative F-actin dataset processed using confocal reconstruction, RL-RbN, s^2^ISM and s^2^ISM-RbN. The closed-pinhole confocal image is shown as a reference to compare the sectioning and resolution gain obtained by exploiting the full detector-array information. Standard s^2^ISM improves lateral resolution and optical sectioning but remains affected by the semi-convergent behaviour of RL-like iterative reconstruction, requiring empirical early stopping to avoid noise amplification. By incorporating the RbN variance regularization into the s^2^ISM forward model, s^2^ISM-RbN preserves the sectioning capability of s^2^ISM while stabilizing the reconstruction against noise overfitting. Magnified regions highlight the recovery of fine filamentous structures and the suppression of reconstruction artefacts in the RbN-regularized result. Scale bars: 5 µm and 1 µm (detail).

**Figure 6:**
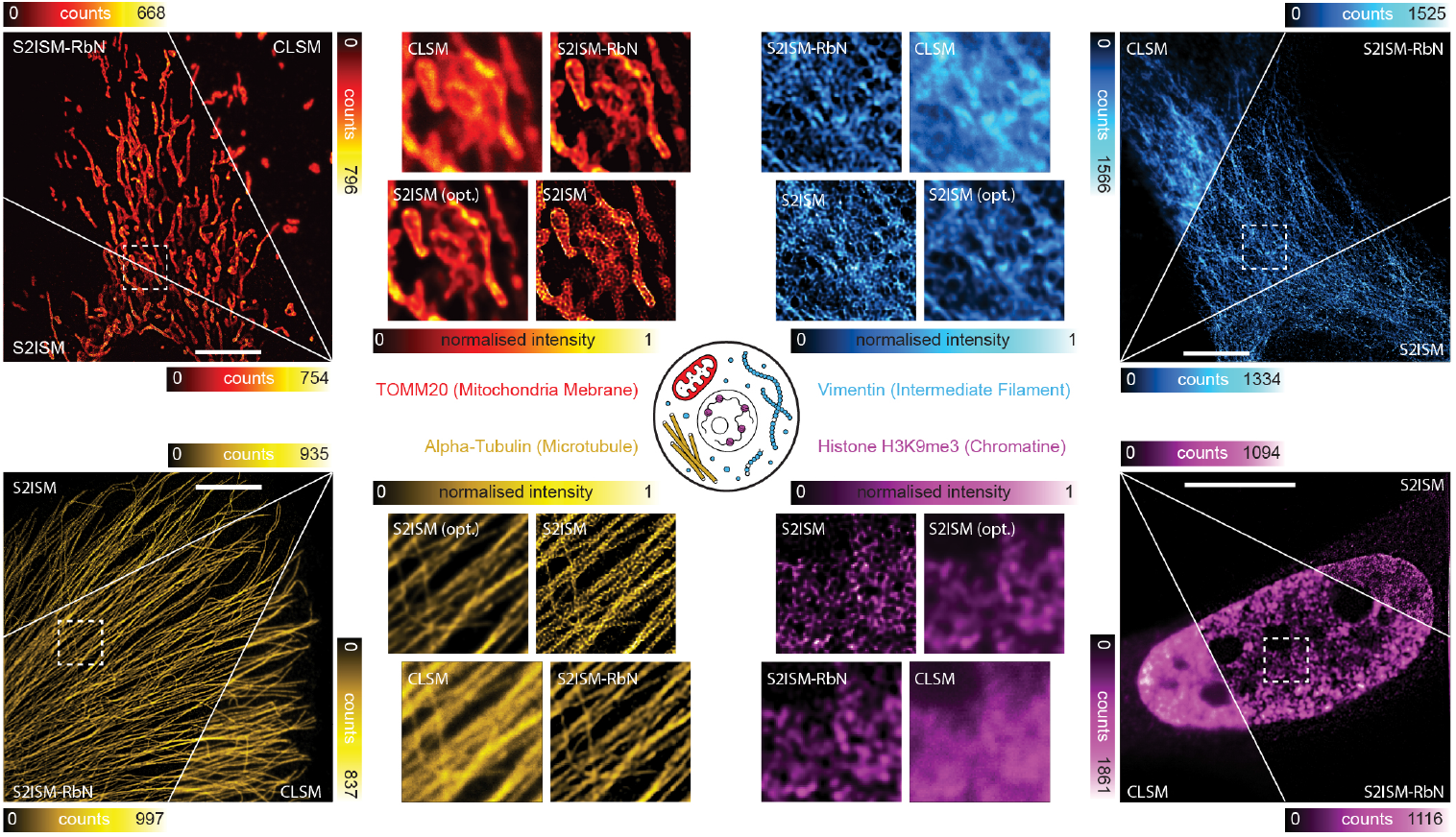
The s^2^ISM-RbN framework was integrated into the s^2^ISM reconstruction pipeline and applied to ISM datasets of the same four subcellular targets shown in Fig. 4 (TOMM20, vimentin, *ε*-tubulin, and histone H3K9me3). For each target, the full field of view is displayed as a diagonal split comparing s^2^ISM and s^2^ISM-RbN, with magnified insets comparing CLSM, s^2^ISM-RbN, s^2^ISM at its empirically optimal stopping iteration [s^2^ISM (opt.)], and over-iterated s^2^ISM. These results demonstrate that s^2^ISM-RbN regularization integrates seamlessly with ISM-based super-resolution reconstruction and generalizes across biological structures and imaging modalities. Scale bars: 5 µm (a).

Together with the results obtained on the custom SPAD-array microscope and on the commercial Evident photon-counting confocal system, these experiments establish RbN as a detector- and forward-model-agnostic regularization framework for photon-counting image reconstruction, rather than a method tied to a specific microscope, acquisition architecture or deconvolution problem.

## 3 Discussion

Single-photon-sensitive detectors are rapidly transforming fluorescence microscopy by faithfully preserving photon-counting statistics while providing richer information at the level of individual photon detections, thereby creating new opportunities for image reconstruction. Here, we show that this additional statistical information can itself become a source of regularization. By exploiting the consistency among independent photon-counting realizations provided by these detectors, the regularized by noise (RbN) framework stabilizes Poisson maximum-likelihood reconstruction without imposing assumptions on specimen morphology. Combined with automatic selection of the regularization strength through the Poisson residual whiteness principle, RbN provides a parameter-free, specimen-agnostic reconstruction strategy that eliminates semi-convergence, avoiding the need for the heuristic early stopping required by conventional Richardson–Lucy reconstruction. The robust performance observed across specimens spanning isolated puncta, filamentous networks, membrane systems and densely organized intracellular structures suggests that statistical consistency provides a genuinely morphology-agnostic source of regularization. This contrasts with conventional regularization strategies, whose effectiveness often depends on assumptions about image smoothness, sparsity or other structural priors, making their performance inherently specimen dependent. Although conceptually related to self-supervised methods based on independent observations, including Noise2Noise, Noise2Self, Noise2Void, and recent gradient-consensus optimization approaches, RbN assigns measurement redundancy a fundamentally different role by embedding statistical consistency directly within the optimization functional. RbN therefore remains within the classical framework of variational regularization, where the reconstruction is defined as the minimizer of an explicit fidelity-plus-regularization functional. This provides a transparent interpretation of the regularizer and enables standard mathematical guarantees, including convergence of the optimization algorithm to a minimizer and consistency of the regularized solutions in the vanishing-noise and vanishing-regularization regime.

We first demonstrated RbN using a custom laser-scanning microscope equipped with an asynchronous-readout SPAD array and a photon-timing-resolved acquisition system. On this platform, the photons detected during each pixel dwell time were partitioned into independent temporal windows, providing multiple statistically independent realizations within a single raster scan. This represents a natural hardware implementation of the framework. The applicability of RbN is not restricted to instruments that directly expose multiple temporal realizations. We additionally applied the framework to data acquired with two commercial photon-counting microscopes: a Nikon system equipped with the NSPARC detector for image scanning microscopy and an Evident laser-scanning confocal microscope based on SiPM detection. For these commercial datasets, independent realizations were generated computationally through stochastic photon splitting of the measured photon counts. More generally, independent realizations could also be obtained from large-format SPAD arrays by grouping high-speed image frames. The successful application of RbN to both systems demonstrates that direct hardware access to temporally resolved photon streams or microscopes capable of directly acquiring independent realizations are not required. Statistically equivalent realizations can instead be generated computationally of any photon-counting measurement, extending the framework to virtually any photon-counting microscope, including systems based on qCMOS cameras.

The successful integration of RbN into the more complex s^2^ISM reconstruction further demonstrates that the framework is independent of a specific image formation model. Despite the substantially more sophisticated forward model, the same regularization principle suppresses noise amplification while preserving resolution enhancement and optical sectioning. RbN therefore regularizes the statistical inverse problem rather than a particular reconstruction algorithm. Its successful application to both conventional confocal deconvolution and s^2^ISM reconstruction, using custom and commercial photon-counting microscopes, demonstrates that the framework is independent of a specific forward model, detector technology or acquisition platform. Although demonstrated here for fluorescence microscopy, the same principle could be extended to other photon-counting inverse problems, including positron emission tomography [58], whenever independent measurement realizations are available.

More broadly, we believe that this work illustrates the beginning of a transition in computational microscopy. Historically, even when using single-photon-sensitive detector, image reconstruction has operated on photon-integrated measurements, where the information carried by individual photon detections is compressed through temporal, and in some case spatial (e.g., laser-scanning microscopy with point detector), integration into a single specimen image. Beyond reflecting the limitations of previous detector and data-acquisition architectures, this representation also substantially reduced data storage, transmission bandwidth, and computational requirements, making the intensity-based image the natural interface between acquisition and reconstruction. Advances in both detector technologies and data-acquisition systems are progressively changing this paradigm by enabling the recording of individual photons together with multidimensional signatures, beyond their arrival time, including spatial and spectral information. The present work represents an intermediate step in this transition. Although RbN reconstruction is still performed from images, the photon stream is first partitioned into multiple statistically independent realizations, allowing the reconstruction algorithm to exploit statistical information that would otherwise be discarded by integration. This representation retains more information than a single integrated image, but it still relies on binning, which irreversibly removes the ordering and precise arrival times of photons within each temporal window. Looking forward, we envisage a new generation of reconstruction algorithms operating directly on the native photon stream, without preliminary aggregation into images. Such methods could jointly exploit not only the statistical independence of individual photon detections, as demonstrated here, but also the temporal relationships arising from fluorescence photophysics, including excited-state dynamics, blinking and photon correlations. A similar transition towards event-level inference is already emerging in single-particle tracking [59, 60]. More generally, we anticipate that future computational microscopy will increasingly operate at the native granularity of the detector output, enabling every detected photon to contribute its full multidimensional information to image reconstruction.

Although RbN was formulated here for Poisson photon-counting data, whose rich statistical information is made accessible by modern single-photon-sensitive detectors, its underlying regularization principle is not intrinsically tied to Poisson statistics. Extending the framework to a different noise model requires adapting both the data-fidelity term and, whenever independent observations are not acquired directly, the strategy used to generate statistically independent realizations, while leaving the consistency-based regularization term unchanged. For additive Gaussian noise, the Poisson Kullback–Leibler fidelity could be replaced by a weighted least-squares term, and independent realizations could either be acquired experimentally or generated computationally when the noise covariance is known [61]. Extending the framework to mixed Poisson–Gaussian noise is more challenging and would require new computational approaches for generating statistically independent realizations. More generally, whenever repeated independent measurements can be acquired directly, the RbN principle could be extended to a broad class of statistical inverse problems through an appropriate noise-specific fidelity and consistency model.

## 4 Methods

### 4.1 Regularisation by Noise framework formulation

The reconstruction problem considered in this work is based on the same physical image formation model underlying Richardson–Lucy (RL) deconvolution. Namely, the measured photon-count image is interpreted as a Poisson realization whose mean is given by the forward projection of the unknown fluorophore distribution through the microscope model. The RL algorithm, derived with a maximum-likelihood formulation (Supplementary section S1.1), is attractive because it combines the optical model of the microscope with the correct photon-counting statistics, and therefore seeks the object that is most likely to have generated the measured data under the assumed Poisson noise. Moreover, RL offers two additional desirable properties that we aim to preserve in the proposed framework: it enforces the non-negativity of the reconstructed image and conserves the total photon flux. However, it is affected by the semi-convergent behaviour: continued iterations progressively amplify noise. To mitigate this instability while retaining the same physics-based Poisson fidelity term and its advantages, we introduce an additional regularization term that preserve the morphology-agnostic character of the RL algorithm. Rather than encoding any prior on the object’s shape, promoting smoothness, sparsity, or any prescribed structural model, that potentially limits the ability to accurately recover the wide variety of structures encountered in biological samples, the proposed regularizer exploits the statistical structure of the acquisition noise, inherently available from multiple independent noise realizations of the same measurement. The rationale is the following. If two or more measurements of the same specimen are acquired under identical conditions, they differ from one another only through their independent Poisson noise realization: at every pixel, each measurement is an independent draw from a Poisson distribution with the same underlying mean. Because these draws are similar to one another, an accurate reconstruction obtained from each individual measurement should, in turn, be similar across measurements. Therefore discrepancies between per-measurement reconstructions are attributable to stochastic noise fluctuations rather than to genuine structural differences; while features that are consistently reconstructed across independent realizations are more likely to represent sample structures. This observation naturally motivates a regularization strategy that penalizes reconstruction variability across realizations.

Let 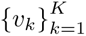 denote *K* independent and identically distributed photon-count realizations associated with the same underlying fluorescence distribution 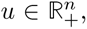 where *n* indicates the number of pixels. Each 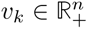 follows

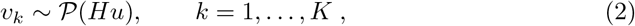

where 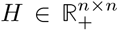 denotes the forward operator encoding the microscope PSF. The RbN formulation translates this statistical redundancy into a regularized variational minimization problem, introducing one auxiliary reconstruction variable *u_k_* for each realization and coupled these variables through a variance penalty

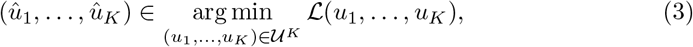

where, denoting by **u** = (*u*_1_*,…, u_K_*) the array of solutions with 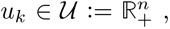 the objective function reads

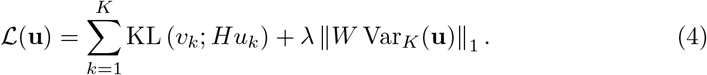

For each pixel *i*, the terms

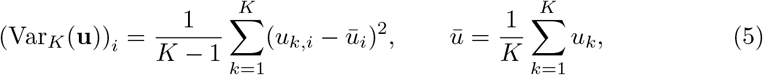

defines the empirical variance image, computed component-wise across the *K* reconstructions. The parameter λ *>* 0 controls the balance between data fidelity and variance regularization, while 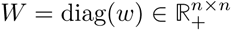 with 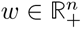 is a diagonal spatial weighting matrix that modulates the contribution of each pixel according to the local reliability of the measurements (see Supplementary section S1.2 for the choice adopted in this work). The final full-flux reconstruction is then obtained as 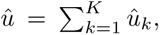 consistently with the fact that 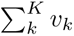 corresponds to the full-dwell-time measurement. The minimization of eq. (3) is performed using an alternating scaled gradient projection scheme that updates the realization-specific variables independently and recomputes their empirical mean, enforcing non-negativity and photon-flux conservation at every iteration. The complete derivation and pseudocode are reported in Supplementary section S1.7 and algorithm S1.

The Poisson data-fidelity term in eq. (4) enforces consistency between the forward model applied to each reconstruction *u_k_*and its corresponding noisy realization *v_k_*, whereas the regularization term penalizes the weighted empirical variance across the ensemble of reconstructed images. Consequently, image features that are consistently recovered across independent realizations incur a small penalty, while features that fluctuate from one realization to another—typically arising from stochastic photon noise—produce a larger variance and are therefore suppressed. Since the regularization acts solely on the agreement among independent reconstructions, rather than on the spatial characteristics of any individual image, it remains agnostic to the morphology of the underlying object. The only prior information exploited by the proposed method is that all measurements are independent realizations of the same object, a property naturally guaranteed by the Poisson photon-counting model and by the acquisition protocol adopted in this work.

### 4.2 Poisson Residual Whiteness Principle

The performance of the proposed RbN framework depends on the choice of the regularization parameter *ω >* 0, which controls the balance between data fidelity and the agreement among independent reconstructions. Differently from the RL iteration counter, this parameter can be interpreted as a user-controllable parameter with a clear optimization meaning: small choices of *ω* give more weight to the data-fidelity term and therefore favour stronger resolution recovery at the cost of noise amplification; while large values of *ω* give more weight to the RbN penalty and favour smoother reconstructions at the cost of lower resolution. In the following, we adapt the Poisson Residual Whiteness Principle (PRWP) [46–48] to the proposed reconstruction model to automatically set the balance between reconstruction and denoising.

The underlying idea of PRWP is that an accurate reconstruction close to the desired unknown ground truth object *û_k_*(λ) ≈ *u_k_*should explain all structured information contained in the measurements. Therefore the residual between the acquired data and the predicted data should contain only statistically uncorrelated Poisson noise. Mathematically, for a given candidate *ω*, the corresponding *û_k_*(*ω*) is mapped into the measurement space through the forward imaging model, yielding the predicted photon-count image *Hû_k_*(*ω*), and then compared to the corresponding raw realization *v_k_*via the standardized residual

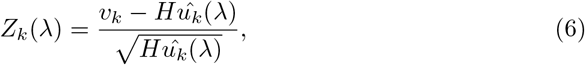

where the square root and division are taken element-wise. Under the assumption that the reconstruction explains well the observed data, *û_k_*(λ) approximate the mean of *v_k_*. Therefore *Z_k_*(λ) should behave as a spatially white random field: zero mean, unit variance, and spatially uncorrelated. The PRWP evaluate the residual above: residuals containing structured image features indicate that the reconstruction has failed to explain part of the measured signal, whereas purely random residuals correspond to a reconstruction consistent with the statistical noise model. To quantify this the autocorrelation of the residual is computed. In details, the PRWP selects the optimal regularization weight as a minimizer of the total residual whiteness measure, computed jointly over the *K* realizations,

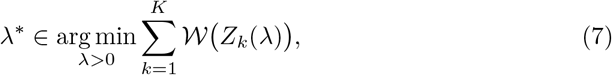

where *W* denotes the squared norm of the autocorrelation. The whiteness (potentially non-convex) function is evaluated over a logarithmically-spaced grid of λ values to find its minimum.

Intuitively, the criterion can be understood as follows: if *ω* is overestimated, the reconstruction remains insufficiently deconvolved and the forward projection fails to reproduce the finest structures observed in the experimental image. The residuals therefore retain spatially correlated features inherited from the sample. On the contrary, if *ω* is underestimated, the reconstruction becomes progressively affected by noise overfitting. Although the subsequent forward projection partially reduces these artefacts through convolution with the PSF, subtle residual correlations remain detectable by the whiteness metric. Looking for minima of the residual autocorrelation thus identify the value of *ω* that best explains the measured data while avoiding both under- and over-regularization.

The whiteness criterion defined above treats every pixel equally. In practice, however, fluorescence microscopy Poisson acquisitions may contain zero-photon pixels (especially in the low-photon count regime). These pixels provide limited information on residual spatial correlations and systematically bias the whiteness functional toward spuriously low values [47, 62], making the parameter selection unstable. To avoid this effect, we adopt a masked version of the Poisson Residual Whiteness Principle proposed in [46, 47] where the analysis gets restricted to the active set of pixels in which at least one photon has been detected Supplementary section S1.3. Within the proposed RbN framework, this criterion provides an automatic, data-driven estimate of the regularization strength without requiring reference images, prior knowledge of the sample, or empirical tuning.

### 4.3 Construction of multi-realization datasets with tunable SNR from a time-resolved SPAD-array acquisition

The proposed RbN framework requires multiple statistically independent realizations of the same underlying fluorescent distribution. The most straightforward experimental strategy would be to acquire a sequence of images of the same specimen under identical imaging conditions. Although this naturally provides independent Poisson realizations, it substantially increases the acquisition time and makes the measurement more susceptible to motion artefacts—precluding its application to dynamic samples such as live cells—and to photobleaching. In this work, independent realizations are instead obtained either computationally, through a stochastic photon-splitting strategy, or experimentally using a dedicated intra-pixel time-resolved acquisition implemented on our SPAD-based microscope. Since photon detections constitute a Poisson point process, photon counts accumulated over disjoint temporal intervals are statistically independent. Consequently, summing photon counts over disjoint subsets of temporal bins yields independent Poisson realizations of the same underlying fluorescent distribution. From time-resolved acquisition, multiple statistically independent sets of noise realizations can be generated at different signal-to-noise ratios.

#### Different SNR

To build the statistically independent noise realizations required by RbN framework at different SNR, the time-resolved tensor 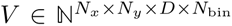 is partitioned along its temporal dimension. Let *n_b_*denote the number of temporal bins assigned to each realization. This defines an equivalent pixel dwell time *dt* = *n_b_ T*_bin_, where *T*_bin_ = *T_dw_/N*_bin_ is the temporal-bin duration and *T_dw_* is the acquisition pixel dwell time. Increasing *n_b_*is therefore equivalent to increasing the effective integration time and consequently the detected photon counts. For a prescribed number *K* of noise realizations and a desired integration factor *n_b_* (with *K n_b_ ≤ N*_bin_)a *K*-realization dataset is built as follows: *K* × *n_b_* temporal bins are randomly sampled without replacement from the available *N_bin_* bins. These sampled bins are arranged into a matrix of size *K* × *n_b_* whose *k*-th row contains the temporal bins assigned to the *k*-th realization. For each *k* = 1*, …, K*, sum the *K* × *n_b_* photon counts matrix over the rows produces *K* independent channel-resolved (or channel-summed, for the confocal case) noise realization at equivalent dwell time *dt* (see Supplementary Fig. S16). Because each temporal bin contributes to at most one realization, the generated datasets remain statistically independent while preserving the Poisson statistics of the photon-counting process. Different SNR regimes are obtained by repeating this procedure with different values of *n_b_*: larger *n_b_* (longer equivalent dwell time) yields higher-SNR realizations; smaller *n_b_* yields lower-SNR realizations. The underlying object and imaging conditions are kept fixed. Throughout this work, three representative conditions were considered by using different values of *n_b_*= 1, 4, 6, corresponding to low-, medium-, and high-SNR respectively acquisitions.

#### Multiple experiments

While the procedure described above yields, for a fixed integration factor *n_b_*, a single set of *K* statistically independent realizations, the specific outcome of this construction depends on which temporal bins are randomly assigned to each realization. To assess the statistical robustness of the reconstruction with respect to this random-partitioning step, and more generally to the specific noise realization drawn from the acquisition, the entire procedure described above is repeated *L* independent times for each SNR regime. At each repetition *l* = 1*, …, L*, a new set of *K* × *n_b_* temporal bins is randomly sampled without replacement from the same time-resolved acquisition, producing a new set of *K* independent noise realizations. The underlying raw acquisition, sample, SNR and microscope configuration were therefore kept fixed, while only the temporal-bin assignment per realization changed. This procedure generates *L* statistically independent multi-realization datasets per SNR regime corresponding to the same biological structure. Since all repetitions originate from the same raw acquisition and differ only in their photon-noise sampling, the variability across the corresponding reconstructions provides a direct estimate of the stability of the reconstruction algorithm to noise.

### 4.4 Computational photon splitting

For photon-counting acquisitions lacking multiple time-resolved realizations, the statistically independent and identically distributed measurements required by RbN can be generated from a single image by binomial photon splitting [37]. Each detected photon was independently assigned to one of two virtual images with probability *p* = 1*/*2. By the Poisson thinning property, if *v* ↓ Poisson(*Hu*), the resulting images satisfy *v*_1_*, v*_2_ ∼ Poisson(*Hu/*2) with *v*_1_ independent from *v*_2_ and *v* = *v*_1_ + *v*_2_. Iterating this splitting procedure yields *K* mutually independent Poisson realizations, *v*_1_*,…, v_K_*, of the same underlying object from a single photon-counting acquisition, making the RbN framework suitable also for non time-resolved acquisition.

### 4.5 Full-reference distortion metrics

For the simulated experiments, where the ground-truth object is known, we evaluated the reconstruction quality using a set of full-reference image-quality metrics (Supplementary Figs. S2 to S4). Let 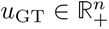 be the ground-truth image with *n* pixels and let 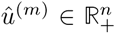 denote the reconstruction at iteration *m*. The root-mean-square error (RMSE) was computed as

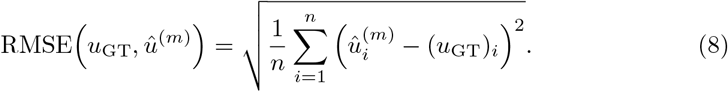

Lower RMSE values indicate smaller pixel-wise deviations from the ground truth. The peak signal-to-noise ratio (PSNR) was computed from the mean-square error after flux normalization of both images,

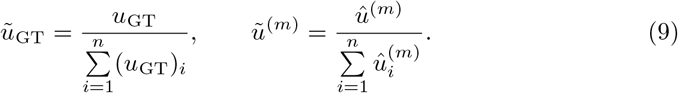

Defining

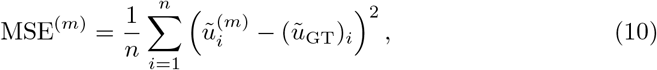

the PSNR was given by

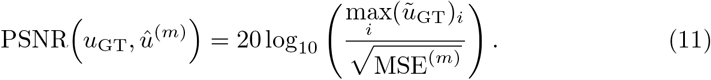

Higher PSNR values indicate a smaller normalized reconstruction error.

The Kullback–Leibler divergence between the normalized ground truth and reconstruction is defined as

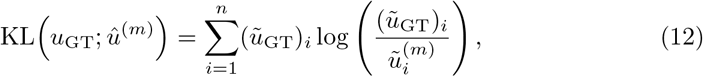

Lower KL values indicate better agreement with the ground-truth intensity distribution. Structural similarity was quantified using the structural similarity index measure (SSIM). For two image patches *x* and *y*, SSIM is defined as

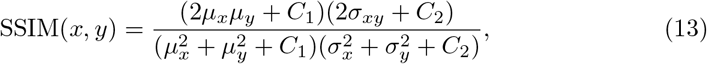

where *µ_x_* and *µ_y_* are the local means, 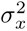 and 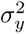 are the local variances, σ*_xy_* is the local covariance, and *C*_1_*, C*_2_ *>* 0 are stabilizing constants. The final SSIM value was obtained by averaging the local SSIM map over the image. Higher SSIM values indicate greater structural similarity to the ground truth.

Because conventional SSIM was originally developed for natural images and can behave unexpectedly on microscopy data with different intensity ranges, background offsets and photon budgets, we additionally computed Micro-SSIM [55]. Micro-SSIM modifies the SSIM comparison by first applying microscopy-specific preprocessing, denoted by *F*(*·*), including background subtraction and range normalization, and then estimating a positive scaling factor *ε >* 0 that maps the prediction intensity range to that of the target image before computing the SSIM score. The Micro-SSIM score can be written as

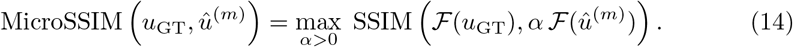

Higher Micro-SSIM values indicate higher microscopy-adapted structural similarity.

### 4.6 Noise level and reconstruction stability

To quantitatively evaluate the robustness of the reconstruction algorithms to measurement noise, we exploited, for each SNR regime, the multiple independently datasets described in Section 4.3, obtained under identical acquisition and reconstruction settings, yielding reconstructed images *û*^(1)^*,…, û*^(*L*)^. Since all reconstructions originate from the same time-resolved acquisition and differ only in the sampled noise realizations, their variability across repetitions provides a direct estimate of the reconstruction stability to noise. Therefore, the pixel-wise standard deviation across repetitions was evaluated as

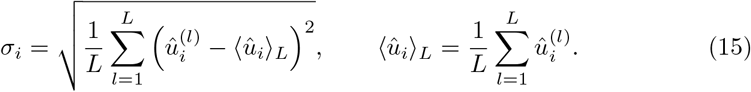

where ⟨·⟩*_L_* denotes the average across the *L* repetitions. Since datasets acquired at different SNRs exhibit different average intensities, direct comparison of the variance is not meaningful. The variance was therefore normalized by the mean reconstructed intensity. The normalized noise-level metric is defined as the pixel-averaged standard deviation normalized by the pixel-averaged mean intensity, namely

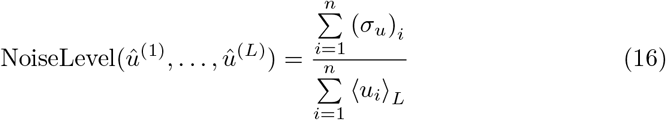

where

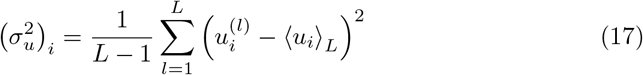

and

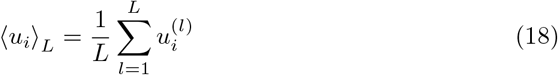

The resulting quantity provides a dimensionless estimate of the reconstruction sensitivity to noise, compensates for the different photon budgets associated with the different SNR regimes enabling a fair comparison of reconstruction stability across conditions. Throughout this work it is referred to as the noise stability metric. Lower values indicate that the reconstruction is less affected by variations of the input noise realization.

### 4.7 Simulated datasets

Synthetic phantoms and corresponding CLSM and ISM point spread functions (PSFs) were generated using the open-source Python framework BrightEyes-ISM [53, 63]. The simulated detection system consisted of a 5 × 5 SPAD array modelled as a matrix of pinhole detectors with a pitch of 75 µm and an active area diameter of 50 µm. All simulations were performed assuming an optical magnification of *M* = 450, an objective numerical aperture of 1.4, and a refractive index of *n* = 1.51, corresponding to standard oil-immersion imaging conditions. Excitation and emission wavelengths were set to 640 nm and 660 nm, respectively. Under these conditions, the detector array corresponded to an effective size of approximately 1.4 Airy Units (A.U.) in the sample plane, while the central detector element corresponded to approximately 0.2 A.U. All synthetic datasets were generated assuming the detector-array acquisition of an ISM system. Open-pinhole CLSM images were obtained by summing the 25 detector images without applying pixel reassignment, whereas closed-pinhole CLSM images were obtained by considering only the central detector element. The corresponding CLSM PSFs were generated following the same procedure by summing the detector-specific ISM PSFs or selecting only the central PSF, respectively. Three different ground-truth phantoms were considered, representing filamentous, extended, and mixed morphologies. Each phantom was convolved with the ISM PSFs to simulate the blurring introduced by the microscope image formation process (Supplementary section S1.1). Finally, Poisson-distributed photon-counting noise was applied to the noiseless images to generate the synthetic measurements.

### 4.8 Fluorescence Microscopes

#### Custom-built LSM with SPAD array detector

Most cellular datasets were acquired using a custom-built laser-scanning microscope (LSM) equipped with a 5 × 5 square single-photon avalanche diode (SPAD) array detector (PRISM Light Kit, Genoa Instruments), previously described in [64, 65]. The system acquires the detector-array data required to generate both confocal laser-scanning microscopy (CLSM) and image scanning microscopy (ISM) datasets. Briefly, the system was designed to image samples labelled with far-red fluorophores by focusing a 640 nm laser diode (PicoQuant LDH-D-C-640S) through a high-numerical-aperture oil-immersion objective lens (Nikon SR HP Apo TIRF 100×/1.49 NA oil). The emitted fluorescence was collected by the same objective, spectrally separated from the excitation light using a dichroic mirror (F48-643, AHF, Germany) and emission filters (LP01-633R-25 and Notch 642), and subsequently imaged onto a secondary image plane where the SPAD array was located. A relay-lens system provided an overall magnification of approximately 500× between the sample plane and the detector plane, corresponding to an effective SPAD-array size of approximately 1 Airy Unit (A.U.). The excitation focus was raster-scanned across the specimen using a pair of galvanometric mirrors. Microscope control, synchronization between scanning and photon detection, and data acquisition were performed using the open-source BrightEyes-MCS software [66].

For every detected photon, the BrightEyes-MCS data-acquisition module records the detector element in which the photon is detected, its arrival time relative to the excitation pulse (when pulsed excitation is used), and its arrival time relative to the pixel clock [67, 68]. Since fluorescence lifetime information was not used in this work, the photon arrival time relative to the excitation pulse was discarded. Instead, the photon arrival time relative to the pixel clock was divided into *N*_bin_ intra-pixel temporal bins spanning the pixel dwell time. Photon events were then accumulated into a four-dimensional photon-counting tensor 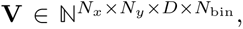 where *N_x_* and *N_y_* denote the raster-scan dimensions, *D* = 25 is the number of SPAD detector elements, and *N*_bin_ is the number of temporal bins within each pixel dwell time. Each tensor element therefore stores the number of photons detected at a given scan position, by a given detector element, and within a given temporal interval of the corresponding pixel dwell time. Conventional CLSM datasets were obtained either by summing over the detector dimension, yielding an open-pinhole image (1 A.U.), or by selecting only the central detector element, corresponding to a closed-pinhole image (0.2 A.U.). The full detector-array data were instead used for ISM reconstruction. The temporal partitioning of the photon stream enables the generation of multiple statistically independent realizations for both CLSM and ISM datasets. The acquisitions are performed with a pixel dwell time of 64 *µ*s (64 time bins of 1 *µ*s), a pixel size of 50 nm and a laser power of 40 *µ*W. The excitation power was chosen to ensure operation below the saturation limit of the detection system, thereby preserving the Poisson statistics of the measurements.

#### Nikon AX with NSPARC detector

Tissue imaging was performed using a commercial Nikon AX laser-scanning confocal microscope built on an inverted Ti2 stand and equipped with an NSPARC spatial-array detector. Both datasets were acquired using a Plan Apo 60× oil-immersion objective with a numerical aperture of 1.42 and an immersion-medium refractive index of 1.515. Images were recorded with the NSPARC detector, a 5 × 5 asynchronous readout SPAD array detector working exclusively in photon-counting mode and providing the raw dataset for ISM reconstruction. Images were recorded using an unidirectional galvanometric scanning, a pixel dwell time of 1 µs, and a 1024 × 1024-pixel raster. The calibrated lateral pixel size was 68.5 nm, corresponding to a field of view of approximately 70.15 × 70.15 µm^2^. The NSPARC projection zoom, corresponding to the virtual detection aperture, was set to 1.0 Airy unit. For imaging EGFP-expressing Purkinje cells in the Thy1-EGFP mouse cerebellum slice, fluorescence was excited at 488 nm and detected over the 502–546 nm spectral range. For the mouse kidney dataset, fluorescence was excited at 561 nm and detected over the 553–618 nm spectral range. Since the acquisition did not provide the intra-pixel temporal partition available on our custom SPAD-baseed LSM system, the statistically independent realizations required by RbN were generated computationally through stochastic photon splitting (Section 4.4).

#### Evident FV5000 with SilVIR detector

Additional confocal datasets were acquired using a commercial Evident FV5000 laser-scanning microscope built on an inverted IX83 platform and operated in photon-counting mode thanks to the SilVIR detector module which integrate a SiPM. The TOMM20 dataset was acquired using a UPLXAPO 60× oil-immersion objective with a numerical aperture of 1.42, an immersion-medium refractive index of 1.518 and a working distance of 0.15 mm. Alexa Fluor 647 was excited at 640 nm, and fluorescence was collected over the 650–710 nm spectral range. Images were recorded using galvanometric scanning with a scan zoom of 1.9×, a nominal pixel dwell time of 2 µs, twofold line integration. The confocal pinhole was operated in automatic mode with an active diameter of 263 µm, corresponding to the nominal 1.0 Airy unit acquisition setting. Since each acquisition provided a single photon-integrated image, the statistically independent realizations required by RbN were generated computationally through stochastic photon splitting and reconstructed using the conventional confocal forward model.

### 4.9 Biological sample preparation

#### Cell Culture

HeLa cells were grown in DMEM with 10% FBS, 1% L-glutamin and 1% penicillin/streptomycin (Sigma-Aldrich) at 37°C and 5% CO2 in a cell culture incubator. After two days in incubator, they were seeded at medium confluence on cleaned and round 18 mm diameter high resolution 1.5” glass coverslips (Marienfield, VWR).

#### Cell fixation

After 24 hours, the cells were washed one time with sterile Dulbecco PBS 1X (Sigma Life Science). A fixation solution (0.2% glutaraldehyde + 0.2% formaldehyde in DPBS) is added for 15 min at room temperature. Cells were then washed 3 times in PBS 1x and stored in PBS 1x + sodium azide 0.1% at 4°C until labelling.

#### Cell immunolabeling

For the saturation step, cells are blocked for 30 min in PBS + 3% BSA + 0.1% Triton. The cells were incubated 1h at room temperature with primary antibody in PBS + 3% BSA + 0.1% Triton. This was followed by three washing steps in PBS + 3% BSA + 0.1% Triton and incubation for 1h at room temperature with secondary antibody. Three more washes with PBS 1x are performed. The cells are then embedded in Prolong Diamond mounting medium (ThermoFisher) and sealed with two-component sealants. Samples are stored in fridge at 4*^↔^*C until imaging.

#### Primary antibodies

The primary antibodies are used with a dilution of 1:500. Mouse anti-alpha-tubulin (Thermo Fisher DM1A + Thermo Fisher B512), rabbit anti-clathrin heavy chain (Abcam ab21679), rabbit anti-TOMM20 (Abcam ab232589), chicken anti-vimentin (EnCor CPCA-Vim), rabbit anti-H3K9me3 (Abcam ab8898), rabbit anti-lamin B1 (ab16048), and rabbit anti-Nup-153 (Abcam, ab84872).

#### Secondary antibodies

The secondary antibodies are used with a dilution of 1:500. Anti-mouse Atto 647N (Sigma 50185), anti-rabbit Alexa Fluor 647 (Thermo Fisher A21246), anti-rabbit Atto 647N (Sigma 40839), anti-chicken CF660C (Biotium 20371), and anti-rabbit abberior STAR 635P (abberior, 1:1,000). F-actin is stained with phalloidin Alexa Fluor Plus 647 (Thermo Fisher A30107).

#### Mouse kidney tissue

The FluoCells prepared slide #3 from Invitrogen. The slides contains a 16 *µ* cryostat section of mouse kidney stained with Alexa Fluor 488 wheat germ agglutinin, Alexa Fluor 568 phalloidin and DAPI.

#### Cerebellum slice

Brains from Thy1-EGFP transgenic mice were fixed overnight at 4,° C in 4% (w/v) paraformaldehyde in phosphate-buffered saline. The tissue was embedded in 2% agarose and sectioned into 300–350*, µ*m slices using a Leica vibrating microtome. Sections were permeabilized for 24 h at 35,° C in 2% (v/v) Triton X-100 in PBS. The tissue was subsequently cleared using RapiClear 1.52 according to the manufacturer’s instructions and mounted using an iSpacer. Purkinje cells were visualized through endogenous EGFP expression in the Thy1-EGFP line.

## 4.10 Data and code availability

All data and code used in this study will be made publicly available through a public repository upon publication of the associated peer-reviewed article.

## Acknowledgements

The work of L.C. is partially supported by the Teaching Science to Computer Flagship at the Italian Institute of Technology. The work of S.A. and Lu.C. is supported by the funding received from the European Research Council (ERC) Starting project MALIN under the European Union’s Horizon Europe programme (grant agreement No. 101117133). The authors thank Dr. Mattia Pesce (Neurofacility, Istituto Italiano di Tecnologia) for acquiring the Evident FluoView datasets, Marco Scotto (Molecular Microscopy and Spectroscopy, Istituto Italiano di Tecnologia) for acquiring the Nikon AX datasets, and Genoa Instruments for technical support with the PrismLight kit.

## Author contributions

A.Z. and G.V. conceived the Regularized by Noise (RbN) framework. Li.C., A.Z., S.S. and Lu.C. derived the RbN optimisation algorithm. Li.C., S.A. and Lu.C. extended the Poisson residual whiteness principle to the proposed regularization framework. Li.C. and S.A. developed the Python software implementing RbN. A.Z., L.L., Lu.C. and G.V. designed the experimental validation. L.L. acquired the microscopy datasets. Lu.C., M.P. and G.V. supervised the project. Li.C., Lu.C. and G.V. analysed and interpreted the results. Li.C., Lu.C. and G.V. wrote the manuscript with input from all authors. All authors discussed the results and approved the manuscript.

