## Supplementary Notes for "Independent noise realizations enable morphologically agnostic image reconstruction in single-photon-sensitive microscopy"

### S1 Supplementary Notes

#### S1.1 Image formation model and the Richardson–Lucy algorithm

In fluorescence microscopy, the image formation under incoherent illumination is modeled as as a spatially invariant, linear blurring process followed by photon-counting (Poisson) detection. Let $u \in \mathbb{R}_+^n$ denote the unknown discretized fluorophore distribution of the object over $n$ pixels, and let $H$ represent the matrix encoding the discretized forward operator, which models convolution with the microscope point spread function (PSF) denoted as $h$. The expected photon-count image is obtained by convolution of the object $u$ with $h$, and the measured image is modelled as a pixel-wise independent realization of a Poisson random process

$$v \sim \mathcal{P}(h * u) = \mathcal{P}(Hu) \ . \tag{S1}$$

The inverse problem consists in estimating the object $u$ from the noisy observation $v$, given the forward operator $H$. Since it is ill-conditioned owing to the finite bandwidth of the optical transfer function, defined as the Fourier transform of the PSF, direct inversion is unstable and requires dedicated reconstruction algorithms. Under the assumption of Poisson statistics, maximum-likelihood estimation naturally leads to Richardson–Lucy (RL) deconvolution algorithm, which has become one of the most widely adopted iterative reconstruction methods in fluorescence microscopy owing to its statistical consistency and positivity-preserving properties.

Under the Poisson model (S1), the negative log-likelihood of the data $v$ given the object $u$ reads, up to an additive constant independent of $u$,

$$-\log \mathbb{P}(v \mid u) \ = \ \sum_{i=1}^{n} \Big[ (Hu)_i - v_i \log(Hu)_i \Big] + \text{const} \ = \ \text{KL}(v; Hu) + \text{const} \ . \tag{S2}$$

Here the KL function indicates the Kullback–Leibler divergence between two vectors $\mathbf{x}, \mathbf{y} \in \mathbb{R}_+^n$ defined as $\text{KL} : \mathbb{R}_+^n \times \mathbb{R}_+^n \to \mathbb{R}_+$, $\text{KL}(\mathbf{x}; \mathbf{y}) := \sum_i^n \text{kl}(x_i, y_i)$ where $\text{kl} : \mathbb{R}_+ \times \mathbb{R}_+ \to \mathbb{R}_+$ is defined as $\text{kl}(a, b) := b - a + a \log(\frac{a}{b}) = b - a \log(b) + a \log(a) - a$. Maximum-likelihood estimation is therefore equivalent to the constrained minimization problem

$$\hat{u} \in \arg \min_{u \in \mathbb{R}_+^n} \text{KL}(v; Hu) \ . \tag{S3}$$

Problem (S3) is solved by the Richardson-Lucy (RL) algorithm [17]. It is an iterative Expectation-Maximization (EM) scheme designed to minimize this objective function while naturally ensuring the non-negativity constraint ($u \geq 0$). The multiplicative update rule at iteration $m$ is formulated as:

$$u^{(m+1)} = u^{(m)} \odot H^\top \left( \frac{v}{Hu^{(m)}} \right) \ , \tag{S4}$$

where $\odot$ and the fraction denote element-wise product and division, $H^\top$ is the adjoint (back-projection) operator. Iteration (S4) is non-negativity preserving, non-increasing in $KL(v; Hu^{(m)})$, and converges to a maximum-likelihood estimate, but it is well known to be *ill-convergent*: prolonged iteration amplifies high-frequency Poisson noise (semi-convergence). Consequently, conventional RL relies on early stopping as its only practical form of regularization. The optimal stopping iteration, however, depends on the sample morphology, the imaging system, and the measurement SNR, and therefore cannot be determined a priori. Stopping the algorithm prematurely leads to incomplete deconvolution, whereas excessive iterations produce reconstruction artefacts caused by noise overfitting. The Regularized by Noise (RbN) framework proposed in this work removes this trade-off by replacing implicit regularization through early stopping with an explicit regularization based on the statistical consistency of multiple independent noise realizations of the same measurement.

#### S1.2 Noise-adaptive regularisation weighting

The expected variability among independent reconstructions naturally depends on the local signal-to-noise ratio (SNR) of the measurements. In regions with high photon counts, independent realizations exhibit relatively small statistical fluctuations and are therefore expected to yield highly consistent reconstructions. Conversely, in low-photon regions, larger discrepancies among reconstructions arise naturally from Poisson noise. Enforcing the same degree of agreement across all image regions would therefore introduce an unnecessary bias in areas where the measurements are intrinsically less reliable. To account for this effect, the variance regularization in eq. (4) is spatially weighted according to the estimated local SNR of the acquired data. Specifically, we define a diagonal weighting matrix $W \in \mathbb{R}_+^{n\times n}$ whose diagonal entries are inversely proportional to the empirical pixel-wise SNR computed from the $K$ independent noise realizations. Since the variability of the reconstructed images is ultimately driven by the uncertainty in the underlying photon measurements, the local measurement SNR provides a natural estimate of the expected variability of the reconstructions. Mathematically, let

$$\mu_i = \frac{1}{K}\sum_{k=1}^{K} v_{k,i}, \qquad \sigma_i = \sqrt{\frac{1}{K-1}\sum_{k=1}^{K}(v_{k,i}-\mu_i)^2}\,, \tag{S5}$$

denote the empirical mean and standard deviation of the photon counts at pixel $i$. The corresponding local SNR and diagonal entries of $W$ are then defined as

$$\mathrm{SNR}_i = \frac{\mu_i}{\sigma_i}\,, \qquad W_{i,i} = \frac{1}{\mathrm{SNR}_i} = \frac{\sigma_i}{\mu_i} \tag{S6}$$

for $i = 1, \ldots, n$.

Because the regularization term in (4) is given by $\|W\,\mathrm{Var}_K(\mathbf{u})\|_1$, pixels with low SNR (larger entries of $W$) contribute more strongly to the penalty, whereas pixels with high SNR contribute less. Consequently, the variance regularization adapts automatically to the local reliability of the measurements, encouraging stronger agreement among independent reconstructions where photon noise is more pronounced while avoiding unnecessary penalization in regions supported by a high photon budget. The weighting matrix $W$ is computed once from the raw measurements and kept fixed throughout the optimization, whereas the scalar parameter $\lambda > 0$ controls the overall strength of the variance regularization relative to the data-fidelity term.

This adaptive weighting provides two complementary advantages. First, it preserves fine structural details in high-SNR regions, where independent reconstructions are already naturally consistent. Second, it effectively suppresses reconstruction instabilities in low-SNR regions, where the observed variability is predominantly explained by Poisson fluctuations rather than genuine structural differences. Unlike conventional morphology-based regularization strategies, the proposed weighting does not rely on assumptions regarding the spatial structure of the object. Instead, it is entirely determined by the statistical properties of the acquired measurements and therefore adapts automatically to different specimens and acquisition conditions without requiring morphology-dependent tuning.

#### S1.3 Masked Poisson Residual Whiteness Principle

The standard Poisson residual whiteness criterion may become unreliable in low-photon-count acquisitions, where the presence of many zero-valued pixels biases the whiteness functional toward artificially low values. Here, we provide the complete masked formulation adopted in this work, following [46, 47]. For each realization $k$, the whiteness analysis is restricted to the active set of pixels in which at least one photon has been detected, defined as

$$\mathcal{M}_k = \{i \in \{1, \dots, n\} \ : \ (v_k)_i > 0\} . \tag{S7}$$

Conditioning on a strictly positive count changes the underlying statistics: on the active set $\mathcal{M}_k$, the measurements are no longer described by a standard Poisson distribution, but by a zero-truncated Poisson distribution. Therefore, the residuals must be standardized using the conditional mean and variance of this zero-truncated model. For each pixel $i \in \mathcal{M}_k$, these quantities are given by

$$\mu^{+}_{k,i}(\lambda) = \frac{\nu_{k,i}(\lambda)}{1 - \exp\left(-\nu_{k,i}(\lambda)\right)}, \qquad \left(\sigma^{+}_{k,i}(\lambda)\right)^2 = \mu^{+}_{k,i}(\lambda) \left[1 - \mu^{+}_{k,i}(\lambda) \exp(-\nu^{\lambda}_{k,i})\right] , \tag{S8}$$

where $\nu_k(\lambda) = H\hat{u}_k(\lambda)$ denotes the forward mapping of the reconstructed image as a function of the regularization parameter. The masked standardized residual associated with realization $k$ is then defined as

$$Z^{+}_{k,i}(\lambda) = \begin{cases} \dfrac{(v_k)_i - \mu^{+}_{k,i}(\lambda)}{\sigma^{+}_{k,i}(\lambda)} \ , & i \in \mathcal{M}_k, \\ 0 \ , & i \notin \mathcal{M}_k. \end{cases} \tag{S9}$$

Similarly as in the main Methods (Section 4.2), the optimal regularization parameter is finally selected by minimizing the whiteness of the masked residuals jointly over the

$K$ independent realizations,

$$\lambda^* \in \arg\min_{\lambda>0} \sum_{k=1}^{K} \mathcal{W}\big(Z_k^+(\lambda)\big) \,. \tag{S10}$$

#### S1.4 Volumetric image formation model and s$^2$ISM algorithm

Image Scanning Microscopy (ISM) extends conventional laser-scanning confocal microscopy by replacing the single-point detector with a detector array. During raster scanning, all detector elements simultaneously record photons emitted from the same excitation position, thereby acquiring multiple spatially shifted observations of the same fluorescent object. Each detector element acts as an individual confocal detector and is characterized by a distinct effective point spread function (PSF). Specifically, a focused excitation beam is scanned across the sample $u$ at positions $\mathbf{x}_s$, and the emitted fluorescence $v$ is imaged onto a $D$-element detector array

$$v(\mathbf{x}_s \mid \mathbf{x}_d) \;\sim\; \mathcal{P}\left(h(\mathbf{x}_s \mid \mathbf{x}_d) * u(\mathbf{x}_s)\right) \tag{S11}$$

where $*$ denotes the convolution on $\mathbf{x}_s$, $\mathcal{P}$ indicates the Poisson distribution describing the noise, and the corresponding effective PSF is defined as

$$h(\mathbf{x}_s \mid \mathbf{x}_d) = h_{\mathrm{exc}}(-\mathbf{x}_s) \cdot \big[h_{\mathrm{em}}(\mathbf{x}_s) * p(\mathbf{x}_s - \mathbf{x}_d)\big], \tag{S12}$$

where $h_{\mathrm{exc}}$ and $h_{\mathrm{em}}$ are the excitation and emission PSFs, and $p$ describes the detector element's active area.

In [53], this model is extended to account for the three-dimensional distribution of specimens by representing the fluorophore distribution as a stack of $N_z$ discrete axial planes, $u(\mathbf{x}_s, z)$. The effective PSF therefore depends not only on the detector position but also on the axial coordinate, yielding a detector- and plane-dependent response $h(\mathbf{x}_s, z \mid \mathbf{x}_d)$. Because in-focus fluorescence is preferentially concentrated on the central detector elements, whereas out-of-focus fluorescence is distributed more uniformly across the array, the simultaneously acquired detector-resolved measurements contain complementary information about the axial origin of the signal. This information can be exploited to jointly recover lateral super-resolution and optical sectioning. Therefore, the signal recorded by each detector channel is modelled as the Poisson-corrupted superposition of the contributions from all axial planes

$$v(\mathbf{x}_s \mid \mathbf{x}_d) \;\sim\; \mathcal{P}\left(\sum_{z=1}^{N_z} h(\mathbf{x}_s, z \mid \mathbf{x}_d) * u(\mathbf{x}_s, z)\right), \qquad d = 1, \ldots, D. \tag{S13}$$

Denoting by $H_{d,z}$ the circulant convolution matrix for channel $d$ and plane $z$, and by $v_d$, $u_z$ the vectorized channel image and plane object, the eq. (S13) can be rewritten in a matrix form as

$$v_d \sim \mathcal{P}\left(\sum_{z=1}^{N_z} H_{d,z}\, u_z\right), \qquad d = 1, \dots, D. \tag{S14}$$

Introducing the stacked object and measurements

$$\mathbf{u} = \begin{bmatrix} u_1 \\ u_2 \\ \vdots \\ u_{N_z} \end{bmatrix}, \qquad \mathbf{v} = \begin{bmatrix} v_1 \\ v_2 \\ \vdots \\ v_D \end{bmatrix}, \tag{S15}$$

and the corresponding forward operator

$$H = \begin{bmatrix} H_{1,1} & \dots & H_{1,N_z} \\ \vdots & \dots & \ddots \\ H_{D,1} & \dots & H_{D,N_z} \end{bmatrix}, \tag{S16}$$

the acquisition process can again be written in the compact form

$$\mathbf{v} \sim \mathcal{P}\left(H\, \mathbf{u}\right) \tag{S17}$$

This model reduces to the purely lateral ISM model (eq. (S12)) when a single in-focus plane is considered ($N_z = 1$), and provides the additional degrees of freedom, encoded in the channel-and-defocus dependence of $h(\mathbf{x}_s, z \mid \mathbf{x}_d)$, that allow the joint recovery of lateral super-resolution and axial (optical-sectioning) information.

The maximum-likelihood estimate of the axial stack $\{u_z\}_{z=1}^{N_z}$ under model (S14) is obtained by minimizing the sum of channel-wise KL divergences between the measured channel images and their multi-plane forward projection,

$$\{\hat{u}_z\}_{z=1}^{N_z} = \underset{\substack{u_z \in \mathbb{R}^n_+ \\ z=1,\dots,N_z}}{\arg\min} \sum_{d=1}^{D} \mathrm{KL}\left(v_d\,;\, \sum_{z=1}^{N_z} H_{d,z}\, u_z\right). \tag{S18}$$

Problem (S18) generalizes the multi-channel RL-type update to the multi-plane setting, and is solved by the multiplicative, EM-derived iteration

$$u_z^{(m+1)} = u_z^{(m)} \odot \sum_{d=1}^{D} H_{d,z}^{\top} \left( \frac{v_d}{\sum\limits_{z'=1}^{N_z} H_{d,z'} u_{z'}^{(m)}} \right), \qquad z = 1, \dots, N_z \tag{S19}$$

The resulting reconstruction simultaneously estimates the fluorophore distribution in all axial planes while naturally accounting for the channel-dependent ISM image formation process. By explicitly modeling the contribution of out-of-focus fluorescence, s$^2$ISM achieves optical sectioning in addition to the lateral resolution enhancement provided by conventional ISM.

The PSFs used in the confocal and volumetric forward models were computed from a theoretical vectorial diffraction model [66]. The model provides the excitation and detection PSFs for each detector element and for each axial plane entering $H_{d,z}$. To match the theoretical PSFs to the experimental acquisition, we estimated dataset-specific calibration parameters directly from the raw detector-array images, following the procedure introduced in [53]. First, the relative shifts between the scanned images acquired by the inner detector elements were measured and fitted to recover the effective microscope magnification, the orientation of the detector array and its rotation with respect to the scan coordinate system. Second, the axial position of the out-of-focus plane was selected from the simulated PSF stack as the defocus maximizing the diversity of the detector response with respect to the focal plane. The resulting calibrated PSF set was then used to construct the operators $H_{d,z}$ employed in the s$^2$ISM and s$^2$ISM-RbN reconstructions. For the confocal RL and RL-RbN reconstructions, the forward operator $H$ was instead obtained by summing the channel-dependent in-focus PSFs over the detector dimension, consistently with the confocal image obtained by summing the detector-channel measurements. Further details regarding the derivation of the forward model, PSF estimation, its physical interpretation, and the corresponding maximum-likelihood formulation are reported in [53].

#### S1.5 Regularized by Noise extension for s$^2$ISM

The proposed RbN framework naturally extends to the three-dimensional reconstruction inverse problem arising in ISM (eq. (S13)). In this setting, the unknown object is represented by a stack of $N_z$ axial sections. We denote by $\mathbf{u}_k := (u_{k,1}, \ldots, u_{k,N_z})$ the three-dimensional volume associated with the $k$-th noise realization, and by $U := (\mathbf{u}_1, \ldots, \mathbf{u}_K)$ the ensemble of the $K$ realization-specific volumes. As in the two-dimensional confocal formulation (eq. (4)), the regularization exploits the agreement among the independently reconstructed volumes by penalizing the empirical variance across realizations, computed independently for each axial section. The $^2$ISM-RbN inverse problem can thus formulated as

$$(\hat{\mathbf{u}}_1, \ldots, \hat{\mathbf{u}}_K) = \underset{U=(\mathbf{u}_1,\ldots,\mathbf{u}_K)}{\arg\min} \sum_{k=1}^{K} \mathrm{KL}(\mathbf{v}_k; H\mathbf{u}_k) + \lambda \left\| W \, \mathrm{Var}_K(U) \right\|_1 . \tag{S20}$$

where $H$ denotes the forward operator introduced in eq. (S16), accounting jointly for the detector-dependent image formation and the axial imaging model. Expanding the Poisson likelihood over the detector channels and the axial planes yields

$$(\hat{\mathbf{u}}_1, \ldots, \hat{\mathbf{u}}_K) = \underset{U=(\mathbf{u}_1,\ldots,\mathbf{u}_K)}{\arg\min} \sum_{k=1}^{K} \sum_{d=1}^{D} \mathrm{KL}\left( v_{k,d}; \sum_{z=1}^{N_z} H_{d,z} u_{k,z} \right) + \lambda \left\| W \, \mathrm{Var}_K(U) \right\|_1 . \tag{S21}$$

The empirical variance is computed independently for each reconstructed axial section. Specifically, for every $z = 1, \ldots, N_z$ and pixel $i = 1, \ldots, n$

$$(\mathrm{Var}_K(U))_{z,i} = \frac{1}{K-1}\sum_{k=1}^{K}(u_{k,z,i} - \bar{u}_{z,i})^2 \quad \text{with} \quad \bar{u}_{z,i} = \frac{1}{K}\sum_{k=1}^{K} u_{k,z,i}\ , \tag{S22}$$

where the variance is computed component-wise over the $K$ reconstructed realizations. The 3D variance is obtained by stacking the variance images of all axial sections, and the weighting matrix $W$ acts independently on each plane. Its diagonal entries are estimated from the local SNR of the channel-integrated measurements, $\tilde{v}_k = \sum_{d=1}^{D} v_{k,d}$, following the procedure described in Supplementary section S1.2. This, combined with the non-negativity of the variance term, shows that the variance in eq. (S21) is indeed equivalent to:

$$\begin{aligned}\|W\,\mathrm{Var}_K(U)\|_1 &= \sum_{i=1}^{n}\sum_{z=1}^{N_z} w_i\,\mathrm{Var}_K(U)_{z,i} = \frac{1}{K-1}\sum_{k=1}^{K}\sum_{i=1}^{n}\sum_{z=1}^{N_z} w_i(u_{k,z,i} - \bar{u}_{z,i})^2 \\ &= \frac{1}{K-1}\sum_{k=1}^{K}\|\mathbf{u}_k - \bar{\mathbf{u}}\|_W^2\end{aligned} \tag{S23}$$

which proves the separability of the regularization term w.r.t. the realizations. The resulting optimization problem is solved using the same numerical strategy (Supplementary section S1.7) adopted for the two-dimensional formulation, replacing the forward and adjoint operators with their three-dimensional counterparts.

This formulation preserves the physical image formation model of $s^2$ISM while introducing a statistical regularization based solely on the agreement among independent reconstructions. Since every realization images the same three-dimensional fluorophore distribution, genuine structures are consistently recovered across the reconstructed volumes, whereas realization-specific fluctuations remain inconsistent and are therefore penalized through the variance term. The regularization acts exclusively along the realization dimension and is independent of both the detector geometry and the axial image formation model, preserving the optical sectioning capability of $s^2$ISM.

#### S1.6 Extension of Poisson Residual Whiteness Principle to $s^2$ISM

The automatic selection of the regularization parameter described in Section 4.2 can be naturally extended to the volumetric reconstruction framework. In this case, however, the reconstructed object is represented by a stack of axial sections, whereas the experimental measurements are detector-resolved two-dimensional images. Consequently, residuals cannot be evaluated directly on individual axial planes, but must be computed after mapping the reconstructed volume through the $s^2$ISM forward model (eq. (S13)). For a given candidate regularization parameter $\lambda$, the reconstruction $\hat{\mathbf{u}}_k(\lambda)$ is first mapped to the detector space according to the s$^2$ISM forward model in eq. (S13), thus obtaining the predicted measurement

$$\hat{\nu}_{k,d}(\lambda) = \sum_{z=1}^{N_z} H_{d,z}\, \hat{u}_{k,z}(\lambda)\ . \tag{S24}$$

The corresponding standardized residual, per realization, is

$$Z_k(\lambda) = \frac{\sum_{d=1}^{D} v_{k,d} - \hat{\nu}_{k,d}(\lambda)}{\sqrt{\sum_{d=1}^{D} \hat{\nu}_{k,d}(\lambda)}}. \tag{S25}$$

where the square root and division are taken element-wise.

As in the two-dimensional formulation, residual spatial structures indicate that part of the measured information has not been correctly reproduced by the reconstruction and therefore reveal an inaccurate choice of the regularization parameter. The regularization parameter is therefore selected by minimizing the residual whiteness jointly over all independent realizations

$$\lambda^* \in \arg\min_{\lambda>0} \sum_{k=1}^{K} \mathcal{W}\big(Z_k(\lambda)\big), \tag{S26}$$

where $\mathcal{W}(\cdot)$ denotes the squared Frobenius norm of the autocorrelation. Namely, for a non-zero matrix $Z \in \mathbb{R}^{m_1 \times m_2}$

$$\mathcal{W}(Z) := \|S(Z)\|_F^2 = \sum_{(l,h)} s_{l,h}(Z)^2. \tag{S27}$$

where $S(Z)$ is the sample normalized autocorrelation whose entries are

$$s_{l,h}(Z) = \frac{1}{\|Z\|^2} \sum_{(i,j)} z_{i,j}\, z_{i+l,\,j+h}. \tag{S28}$$

Under periodic boundary conditions, the autocorrelation $\mathcal{W}(Z)$ is evaluated efficiently from the 2D discrete Fourier transform $\tilde{Z}$ of $Z$ via Parseval's theorem,

$$\mathcal{W}(Z) = \frac{\sum_{(i,j)} |\tilde{z}_{i,j}|^4}{\big(\sum_{(i,j)} |\tilde{z}_{i,j}|^2\big)^2}, \tag{S29}$$

with $O(n \log n)$ computational cost, where $n = m_1 \times m_2$ is the number of pixels.

The same masking correction described in Section 4.2 is applied to the channel-integrated measurements to avoid dealing with zero photon-count pixels. Here, for each realization $k = 1, \dots, K$, the active set is defined as the set of pixels in which at least one photon has been detected over the entire detector

$$\mathcal{M}_k := \Big\{ i \in \{1, \dots, n\} \; : \; \sum_{d=1}^{D} (\hat{v}_{k,d})_i > 0 \Big\}. \tag{S30}$$

Conditioning on strictly positive photon counts changes the underlying statistics of the channel-summed measurement. Since the sum of independent Poisson variables is still Poisson, the channel-integrated measurement is described by a Poisson distribution with mean the sum of the mean. We can therefore apply the same argument of eq. (S8) and eq. (S9) on $\sum_{d=1}^{D} \hat{v}_{k,d}$.

This masked formulation preserves the physical interpretation of the PRWP established in Sec. 4.2 while accounting for the three-dimensional image formation model of s$^2$ISM. The relevant zero/nonzero condition is naturally defined on the channel-summed measurement rather than on individual detector channels. Since the residuals are evaluated after projection through the complete forward operator, the estimated regularization parameter simultaneously reflects the quality of the lateral reconstruction, the axial sectioning, and the agreement with the experimentally acquired detector images.

#### S1.7 Numerical optimization: Scaled Gradient Projection

The introduction of the consistency regularization term prevents the derivation of a closed-form multiplicative update analogous to the Richardson–Lucy algorithm. In fact, the minimization of the regularized functionals in eq. (4) and eq. (S20) cannot be solved via standard multiplicative RL updates. The optimization problem is solved using the Scaled Gradient Projection (SGP) method, a gradient-based optimization algorithm specifically designed for constrained convex optimization problems and already adopted in fluorescence microscopy inverse problems [42]. The objective function in (4) and eq. (S20) couples the realization-specific variables through the empirical mean. To exploit the resulting structure, we introduce the empirical mean $\bar{u}$ as an auxiliary variable and consider the equivalent lifted formulation

$$\operatorname*{arg\,min}_{\substack{(u_1,\dots,u_K)\in\mathcal{U}^K \\ \bar{u}\in\mathcal{U}}} \sum_{k=1}^{K} \mathrm{KL}(v_k; Hu_k) + \frac{\lambda}{K-1} \sum_{k=1}^{K} \|u_k - \bar{u}\|_W^2, \tag{S31}$$

where the minimizer with respect to $\bar{u}$ is given by the empirical mean $\bar{u} = \frac{1}{K}\sum_{k=1}^{K} u_k$. For fixed $\bar{u}$, the objective is separable with respect to the realization-specific variables, thanks to eq. (S23). We therefore adopt an alternating scheme in which the variables $u_k$ are updated independently while keeping $\bar{u}$ fixed, followed by an update of the empirical mean. We then define:

$$\mathcal{L}_k(u_k; \bar{u}) = \mathrm{KL}(v_k; Hu_k) + \frac{\lambda}{K-1} \|u_k - \bar{u}\|_W^2. \tag{S32}$$

which specializes to eq. (3) for the confocal case ($H$ is the single-channel operator) and to eq. (S20) for the $s^2$ISM case ($H$ is the channel-and-plane operator $\{H_{d,z}\}$, applied and adjoint-applied by summing, respectively, over $z$ and over $d$). The positivity constraint is imposed element-wise on every reconstructed image.

***Variable-metric gradient step.***

For each realization $k = 1, \ldots, K$, the gradient of (S32) with respect to $u_k$ is:

$$\nabla_{u_k} \mathcal{L}_k(u_k; \bar{u}) = \mathbf{1} - H^T \left( \frac{v_k}{H u_k} \right) + \frac{2\lambda}{K-1} W \odot (u_k - \bar{u}). \tag{S33}$$

Given a set of initializations $\left\{ u_k^0 \right\}_k \in \mathbb{R}_+^n$, the SGP algorithm updates the variables by moving along the scaled steepest descent direction followed by a projection onto the non-negative orthant $\mathbb{R}_+^n$, so that the iteration reads:

$$u_k^{(m+1)} = \mathscr{P}_+ \left( u_k^{(m)} - \alpha_k^{(m)} (D_k^{(m)})^{-1} \nabla_{u_k} \mathcal{L}_k(u_k; \bar{u}) \right), \qquad m \geq 0. \tag{S34}$$

where $\mathscr{P}_+$ denotes the projection operator setting negative values to zero and guarantees the physical non-negativity of the reconstructed fluorophore distribution, $\alpha_m$ is the step size chosen via an Armijo backtracking line-search to guarantee stable convergence (the LS2 in [69]), and $D_k^{(m)}$ is a diagonal, positive-definite scaling matrix defined for

$$\left( D_k^{(m)} \right)^{-1} = \operatorname{diag} \left( \frac{u_k^{(m)}}{\mathbf{1} + \frac{2\lambda}{K-1} W u_k^{(m)}} \right). \tag{S35}$$

Both the scaling matrix and the step-size are updated at every iteration according to the adaptive strategy described in [42] and [69], reported here for the sake of clarity.

***Step-length control and metric truncation.***

Following [69] and [42], the step length $\alpha_k^{(m)}$ is selected by an Armijo backtracking procedure with parameter $\delta \in (0, 1]$. Denoting by $\tilde{u}_k^{(m+1)}(\alpha_k^{(m)})$ the tentative update computed from $u_k^{(m)}$ using the step length $\alpha_k^{(m)}$, the sufficient decrease condition reads

$$\begin{aligned} \mathcal{L}_k \left( \tilde{u}_k^{(m+1)}(\alpha_k^{(m)}); \bar{u}^{(m)} \right) - \mathcal{L}_k \left( u_k^{(m)}; \bar{u}^{(m)} \right) \leq & \\ \delta\, \alpha_k^{(m)} \Big\langle \tilde{u}_k^{(m+1)}(\alpha_k^{(m)}) - u_k^{(m)}, (D_k^{(m)})^{-1} \nabla_{u_k} \mathcal{L}_k(u_k^{(m)}; \bar{u}^{(m)}) \Big\rangle, & \end{aligned} \tag{S36}$$

which is enforced independently for each realization $k = 1, \ldots, K$. The metric is truncated to a bounded range to guarantee convergence,

$$(D_k^{(m)})^{-1} = \operatorname{diag} \left( \max \left\{ \frac{1}{\gamma^{(m)}},\ \min \left\{ \gamma^{(m)},\ \frac{u_k^{(m)}}{\mathbf{1} + \frac{2\lambda}{K-1} W u_k^{(m)}} \right\} \right\} \right), \tag{S37}$$

where the thresholding parameter ensuring non-degenerate positive definiteness is

$$\gamma^{(m)} = \sqrt{1 + \frac{s_1}{(m+1)^{s_2}}} \tag{S38}$$

with the parameters $s_1$ and $s_2$ are chosen in this work as $s_1 = 1 \times 10^{18}$, $s_2 = 2$ (see, e.g., [45] for such choice). A large $s_1$ accelerates convergence during early iterations, while $s_2 > 1$ ensures $\gamma^{(m)} \to 1$ asymptotically, recovering the standard Euclidean metric. After updating all realization-specific variables and recomputing the empirical mean, flux conservation is enforced by rescaling each realization so that so that $\|u_k^{(m+1)}\|_1 = \|u_k^{(0)}\|_1 = \|v_k\|_1$.

The complete alternating procedure, including metric truncation, Armijo line search, positivity projection, photon-flux normalization and synchronization through the empirical mean, is summarized in algorithm S1.

##### *Stopping criteria*

At each outer iteration $m \geq 1$, the update (S34) followed by a flux normalization procedure is applied in parallel to each realization $k = 1, \ldots, K$. Once all estimates have been updated, the ensemble mean is recomputed as $\bar{u}^{(m+1)} = \frac{1}{K} \sum_k u_k^{m+1}$ and used in the regularization term of the subsequent iteration; this synchronization step is the only point at which the $K$ update branches interact.

Iterations proceed until either of two convergence criteria is satisfied. The first monitors convergence of the reconstructed object: the algorithm stops when the relative difference between consecutive iterates falls below a prescribed tolerance $\tau$ (here $\tau = 10^{-6}$),

$$\frac{\left\|\bar{u}^{(m+1)} - \bar{u}^{(m)}\right\|}{\left\|\bar{u}^{(m)}\right\|} < \tau, \tag{S39}$$

The second is a safeguard against optimizer stagnation: the algorithm also stops if the step sizes $\alpha_k^{(m)}$ fall below a minimum threshold $\alpha_{\min}$ (here $\alpha_{\min} = 10^{-8}$ for every realization $k$,

$$\max_{k=1,\ldots,K} \alpha_k^{(m)} < \alpha_{\min}, \tag{S40}$$

indicating that the line search can no longer find a non-null stepsize for any realization. A maximum number of iterations (here of 2000) was additionally imposed as a safeguard.

**Algorithm S1** Alternating SGP scheme for RbN reconstruction

**Initialize:** $m = 0$, $u_k^{(0)} \in \mathcal{U}$, $\alpha_k^{(0)} = \alpha_0$, and $c_k = \|v_k\|_1$, for $k = 1, \ldots, K$

**Compute:** $\bar{u}^{(0)} = \frac{1}{K} \sum_{k=1}^{K} u_k^{(0)}$

**while** not converged **do**

    **for** $k = 1, \ldots, K$ **do**

        Compute the truncated metric $(D_k^{(m)})^{-1}$ according to (S37)

        Compute

$$g_k^{(m)} = \nabla_{u_k} \mathcal{L}_k(u_k^{(m)}; \bar{u}^{(m)})$$

        Compute the projected point

$$p_k^{(m)} = \mathcal{P}_+ \Big( u_k^{(m)} - (D_k^{(m)})^{-1} g_k^{(m)} \Big)$$

        Select $\alpha_k^{(m)}$ by Armijo backtracking until (S36) is satisfied

        Set

$$\widetilde{u}_k^{(m+1)} = u_k^{(m)} + \alpha_k^{(m)} \big( p_k^{(m)} - u_k^{(m)} \big)$$

        Enforce flux conservation:

$$u_k^{(m+1)} = \frac{c_k}{\|\widetilde{u}_k^{(m+1)}\|_1} \widetilde{u}_k^{(m+1)}$$

    **end for**

    $\bar{u}^{(m+1)} = \frac{1}{K} \sum_{k=1}^{K} u_k^{(m+1)}$

    $m \leftarrow m + 1$

**end while**

### Supplementary Figures

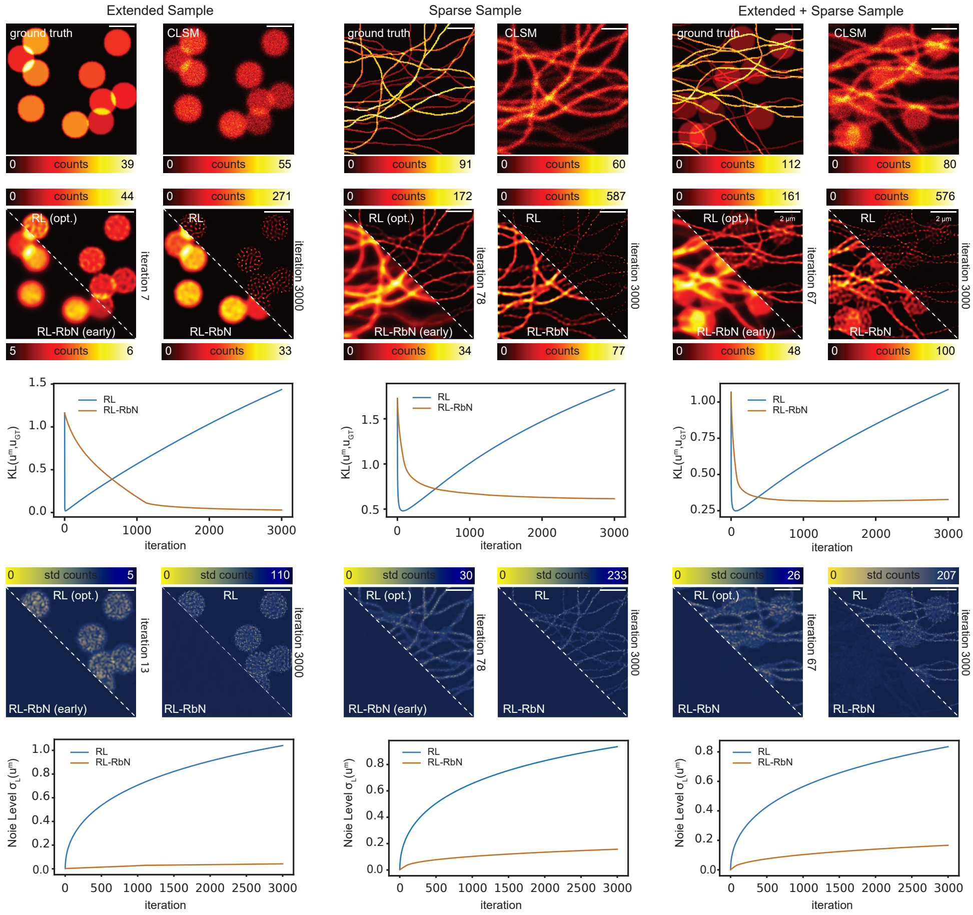


**Figure S1**: **Evaluation of Agnostic Regularization on Simulated Datasets.** Synthetic confocal datasets were generated from three ground-truth objects representing an extended morphology (left), a sparse filamentous morphology (centre), and a mixed morphology containing both extended and sparse structures (right). Synthetic confocal datasets were generated from three ground-truth objects representing an extended morphology (left), a sparse filamentous morphology (centre), and a mixed morphology containing both extended and sparse structures (right). For each object, the ground truth and corresponding noisy CLSM image are shown together with reconstructions obtained using conventional Richardson–Lucy (RL) and regularization by noise (RL-RbN) computed with $K = 8$ noise realizations. RL reconstructions are displayed at the iteration minimizing the Kullback–Leibler divergence from the ground truth, denoted RL (opt.), and after 3000 iterations. RL-RbN reconstructions are shown at an early iteration matched to the RL comparison and at convergence. The Kullback–Leibler divergence, $\mathrm{KL}(\hat{u}^{(m)}, u_{\mathrm{GT}})$, is plotted as a function of iteration number. Reconstruction sensitivity to noise was quantified over $L = 5$ independent noise realizations. Pixel-wise standard-deviation maps are shown for RL and RL-RbN at the indicated iterations, and the corresponding normalized noise level is reported as a function of iteration number. Scale bars, $2\,\mu$m.

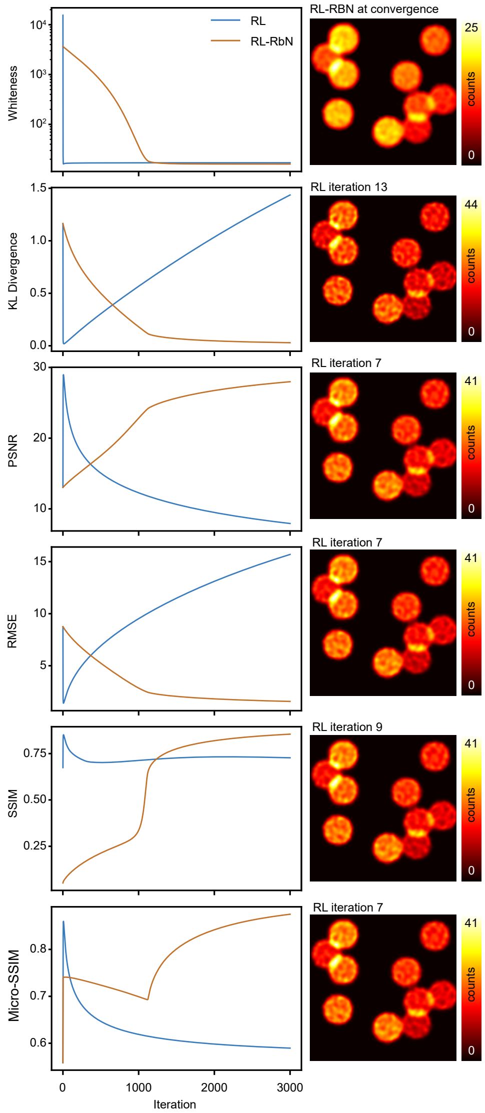


**Figure S2**: **Different image-quality metrics identify different optimal stopping iterations for RL deconvolution of a simulated extended sample.** Evolution of the residual whiteness, Kullback–Leibler divergence, peak signal-to-noise ratio (PSNR), root-mean-square error (RMSE), structural similarity index measure (SSIM), and Micro-SSIM as a function of iteration number for conventional Richardson–Lucy (RL) deconvolution. For metrics describing a reconstruction error, namely KL divergence and RMSE, the optimal reconstruction was defined by the minimum metric value, whereas for PSNR, SSIM, and Micro-SSIM it was defined by the maximum value. The corresponding RL reconstructions are shown on the right at the selected iterations: iteration 13 for KL divergence, iteration 7 for PSNR, iteration 7 for RMSE, iteration 9 for SSIM, and iteration 7 for Micro-SSIM. The RL-RbN reconstruction at convergence is shown as a stable reference.

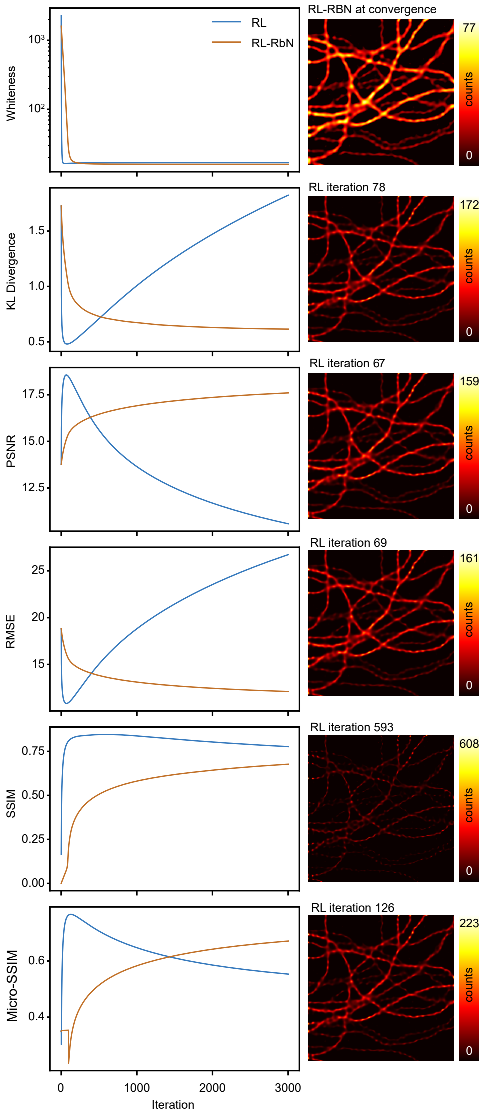


**Figure S3**: **Different image-quality metrics identify substantially different optimal stopping iterations for RL deconvolution of a simulated sparse filamentous sample.** Evolution of the residual whiteness, Kullback–Leibler divergence, peak signal-to-noise ratio (PSNR), root-mean-square error (RMSE), structural similarity index measure (SSIM), and Micro-SSIM as a function of iteration number for conventional Richardson–Lucy (RL) deconvolution and RL-RbN. For metrics describing a reconstruction error, namely KL divergence and RMSE, the optimal reconstruction was defined by the minimum metric value, whereas for PSNR, SSIM, and Micro-SSIM it was defined by the maximum value. The corresponding RL reconstructions are shown on the right at the selected iterations: iteration 78 for KL divergence, iteration 67 for PSNR, iteration 69 for RMSE, iteration 593 for SSIM, and iteration 126 for Micro-SSIM. The RL-RbN reconstruction at convergence is shown as a stable reference.

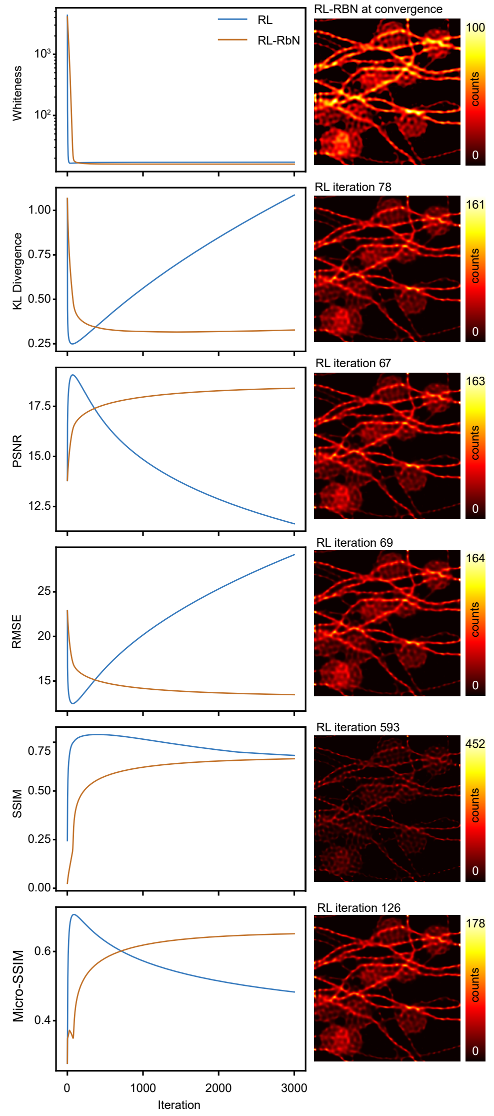


**Figure S4**: **Different image-quality metrics identify substantially different optimal stopping iterations for RL deconvolution of a simulated mixed extended and filamentous sample.** Evolution of the residual whiteness, Kullback–Leibler divergence, peak signal-to-noise ratio (PSNR), root-mean-square error (RMSE), structural similarity index measure (SSIM), and Micro-SSIM as a function of iteration number for conventional Richardson–Lucy (RL) deconvolution and RL-RbN. For metrics describing a reconstruction error, namely KL divergence and RMSE, the optimal reconstruction was defined by the minimum metric value, whereas for PSNR, SSIM, and Micro-SSIM it was defined by the maximum value. The corresponding RL reconstructions are shown on the right at the selected iterations: iteration 78 for KL divergence, iteration 67 for PSNR, iteration 69 for RMSE, iteration 593 for SSIM, and iteration 126 for Micro-SSIM. The RL-RbN reconstruction at convergence is shown as a stable reference.

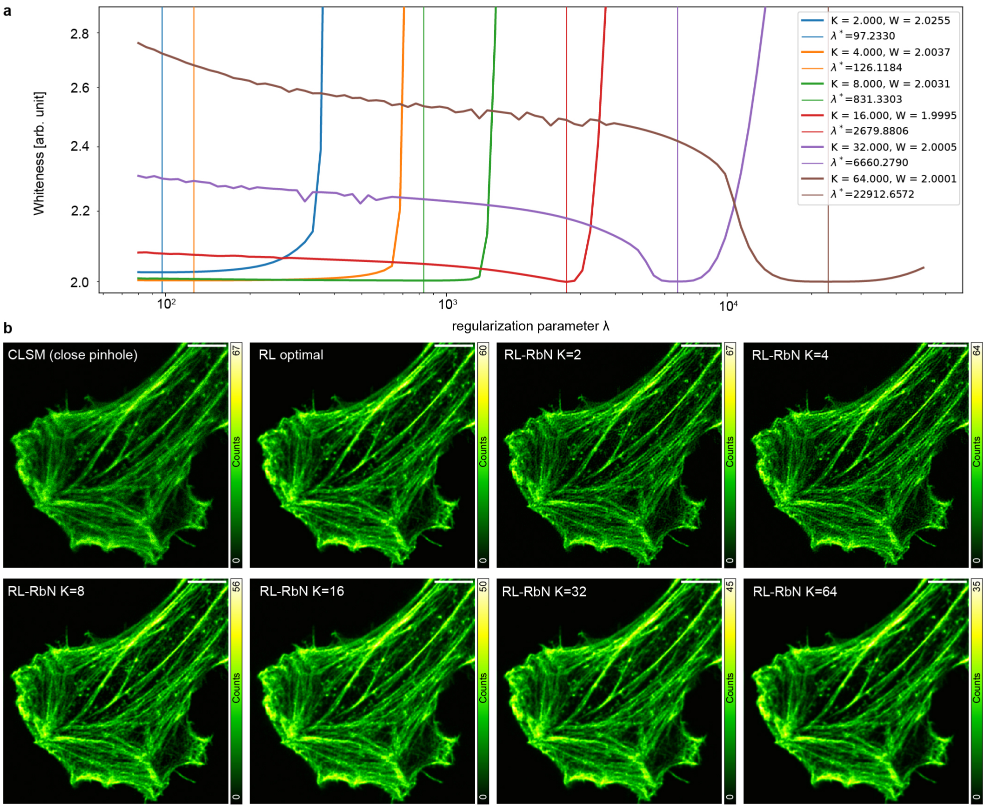


**Figure S5**: **Regularised by noise deconvolution as function of noise realisations number. a,** Residual-whiteness metric as a function of the regularization parameter $\lambda$ for RL-RbN reconstructions obtained using $K = 2, 4, 8, 16, 32$, and 64 independent noise realizations. For each value of $K$, the vertical line indicates the optimum regularization parameter $\lambda^*$ selected by the Poisson residual whiteness principle, whereas the corresponding minimum whiteness value is reported in the legend. The selected regularization strength increases with $K$, reflecting the progressive reduction in photon budget available to each individual realization and the consequent change in the relative balance between the Poisson data-fidelity and variance-regularization terms. **b,** Closed-pinhole CLSM image, optimally early-stopped RL reconstruction, and converged RL-RbN reconstructions obtained for the different values of $K$ using the corresponding automatically selected $\lambda^*$. Increasing $K$ initially improves noise suppression by providing a larger number of independent realizations over which statistical consistency can be enforced. However, excessively large $K$ values reduce the photon counts available in each realization, causing the regularization term to increasingly dominate the data-fidelity term and progressively limiting deconvolution. A value of $K = 8$ therefore provides an effective compromise between statistical redundancy, photon budget per realization, reconstruction quality, and computational complexity, and was used in the subsequent experiments. Scale bars, 5 $\mu$m.

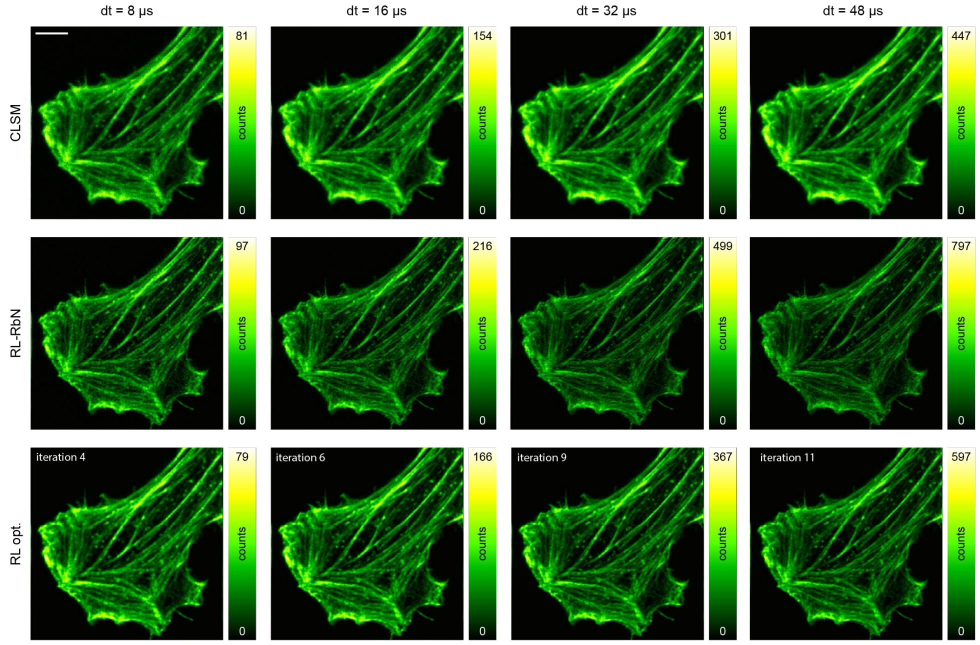


**Figure S6**: **Comparison RL with RL-RbN at different signal-to-noise ratio.** Representative CLSM images of fluorescently labelled F-actin acquired at increasing effective pixel dwell times, $dt = 8$, 16, 32, and 48 $\mu$s, together with the corresponding converged RL-RbN reconstructions and conventional Richardson–Lucy reconstructions stopped at the iteration selected by the Poisson residual whiteness principle, denoted RL (opt.). The automatically selected stopping iterations for RL were 4, 6, 9, and 11 for $dt = 8$, 16, 32, and 48 $\mu$s, respectively. As the photon budget decreases, the optimal RL stopping point shifts towards progressively earlier iterations, limiting the amount of deconvolution that can be performed before noise amplification becomes detectable. Consequently, the difference between RL-RbN and optimally early-stopped RL is most pronounced under photon-limited conditions: RL-RbN preserves a clean background while recovering sharper and more continuous actin filaments, whereas early stopping increasingly restricts structural recovery. At higher photon budgets, RL can be iterated further before the onset of noise overfitting, reducing, but not eliminating, the performance gap between the two methods. Scale bar, 5 $\mu$m.

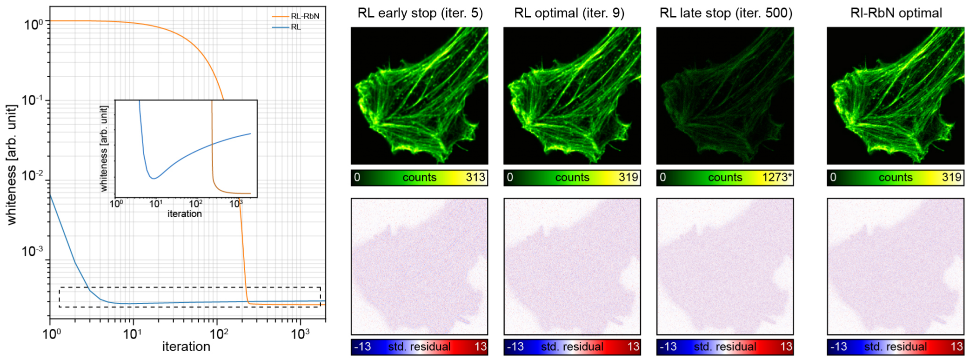


**Figure S7**: **Study of whiteness principle as function of iterations. a,** Residual-whiteness metric as a function of iteration number for conventional Richardson–Lucy (RL) deconvolution and RL-RbN. The dashed box highlights the low-whiteness region reached by the optimally stopped RL reconstruction and by converged RL-RbN. **b,** RL reconstructions obtained before the whiteness minimum [early stop, iteration 5], at the automatically selected optimum [iteration 9], and after prolonged iteration [late stop, iteration 500], compared with the converged RL-RbN reconstruction. The corresponding standardized residuals are shown below. The asterisk (*) indicates that the displayed maximum corresponds to the 99.9th percentile of the pixel-intensity distribution.

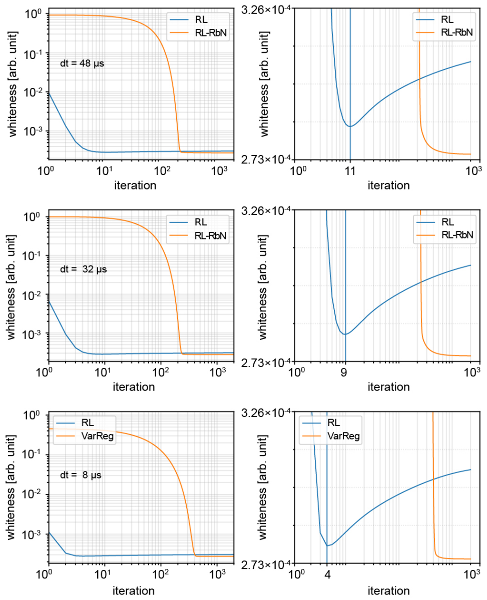


**Figure S8**: **Study of whiteness principle as function of iterations for different SNR.** Residual-whiteness metric as a function of iteration number for conventional Richardson–Lucy (RL) deconvolution and RL-RbN at effective pixel dwell times of $dt = 48$, 32, and $8\,\mu$s, corresponding to progressively decreasing photon budgets. The left column shows the complete evolution of the whiteness metric on logarithmic axes, whereas the right column magnifies the low-whiteness region around the minima. Vertical lines indicate the iteration selected by minimizing the residual whiteness for each reconstruction method. The selected RL stopping iteration shifts from iteration 11 at $dt = 48\,\mu$s to iteration 9 at $dt = 32\,\mu$s and iteration 4 at $dt = 8\,\mu$s. Thus, as the photon budget decreases, RL must be stopped progressively earlier, limiting the amount of recoverable structural information before noise amplification occurs. In contrast, RL-RbN undergoes a longer initial transient before reaching a lower and stable residual-whiteness plateau, consistently avoiding the late-iteration increase observed for conventional RL.

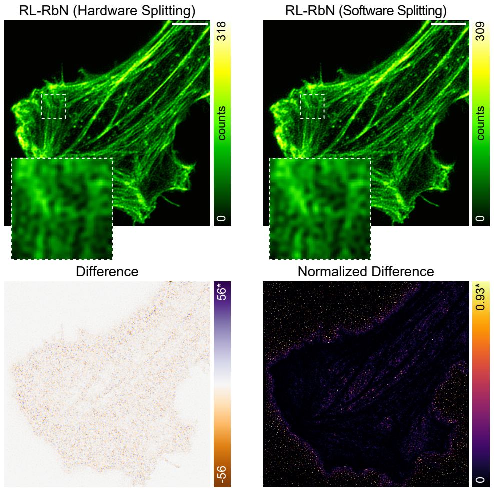


**Figure S9**: **Comparison of computational and hardware photon splitting.** Top: RL-RbN reconstructions of fluorescently labelled F-actin obtained using statistically independent noise realizations generated either experimentally through intra-pixel temporal partitioning of the photon stream (hardware splitting, left) or computationally through stochastic photon splitting of the corresponding photon-counting dataset (software splitting, right). Magnified insets of the same region highlight the close agreement between the two reconstructions in the recovery of fine filamentous structures. Bottom left: pixel-wise difference between the hardware- and software-splitting reconstructions. Bottom right: corresponding absolute normalized difference, obtained after accounting for the overall intensity scale. The asterisk ($^*$) indicates that the displayed maximum corresponds to the 99.9th percentile of the pixel-intensity distribution. Scale bars, 5 $\mu$m.

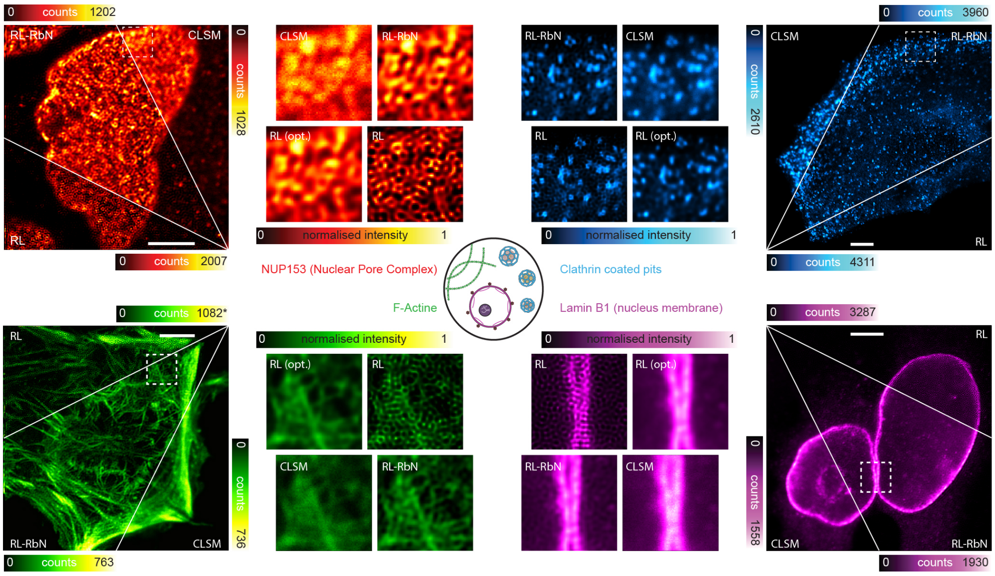


**Figure S10**: **Versatility of RbN on confocal dataset.** RL-RbN was applied to CLSM images of four additional subcellular targets representing markedly different biological organizations: the nuclear pore complex component NUP153, distributed as puncta along the nuclear envelope (red); clathrin-coated pits, appearing as small isolated punctate structures (cyan); F-actin, forming an extended network of intersecting filaments and bundles (green); and lamin B1, delineating the continuous nuclear lamina (magenta). For each target, the full field of view is divided into three triangular sectors showing the raw CLSM image, the RL-RbN reconstruction, and the over-iterated RL reconstruction. Magnified regions compare CLSM, RL-RbN, RL stopped at the PRWP-selected optimal iteration [RL (opt.)], and over-iterated RL. The asterisk (*) indicates that the displayed maximum corresponds to the 99.9th percentile of the pixel-intensity distribution. Scale bars: 5 µm.

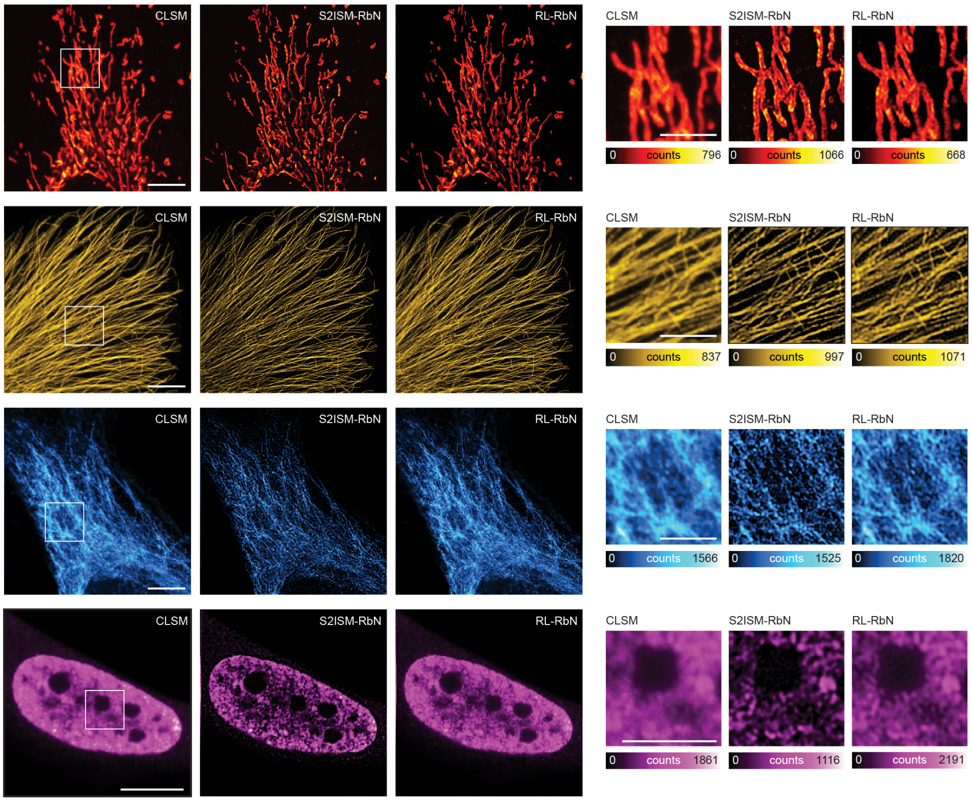


**Figure S11**: **Comparison of CLSM and ISM imaging with our approach.** Representative datasets of four fluorescently labelled subcellular structures with markedly different spatial organizations: TOMM20-labelled mitochondria (red), $\alpha$-tubulin-labelled microtubules (yellow), vimentin intermediate filaments (cyan), and H3K9me3-labelled heterochromatin (magenta). For each specimen, the full field of view is shown as the raw confocal laser-scanning microscopy image (CLSM), the corresponding s$^2$ISM-RbN reconstruction, and the RL-RbN reconstruction obtained from the confocal dataset. The boxed regions are enlarged on the right and displayed for all three reconstruction modalities. Scale bars, 5 $\mu$m for the full fields of view and 1 $\mu$m for the magnified regions.

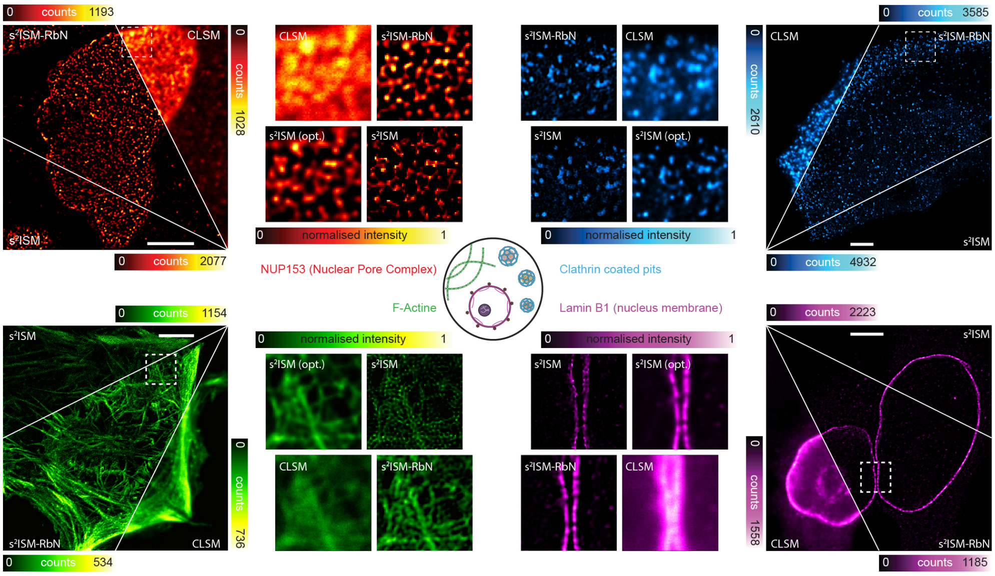


**Figure S12**: **Versatility of RbN on ISM dataset.** s$^2$ISM-RbN was applied to CLSM images of four additional subcellular targets representing markedly different biological organizations: the nuclear pore complex component NUP153, distributed as puncta along the nuclear envelope (red); clathrin-coated pits, appearing as small isolated punctate structures (cyan); F-actin, forming an extended network of intersecting filaments and bundles (green); and lamin B1, delineating the continuous nuclear lamina (magenta). For each target, the full field of view is divided into three triangular sectors showing the raw CLSM image, the s$^2$ISM-RbN reconstruction, and the over-iterated s$^2$ISM reconstruction. Magnified regions compare CLSM, s$^2$ISM-RbN, s$^2$ISM stopped at the PRWP-selected optimal iteration [s$^2$ISM (opt.)], and over-iterated s$^2$ISM. The asterisk (*) indicates that the displayed maximum corresponds to the 99.9th percentile of the pixel-intensity distribution. Scale bars: 5 µm.

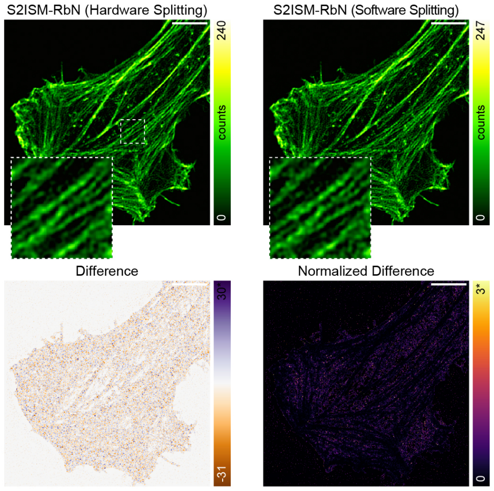


**Figure S13**: **Comparison of computational and hardware photon splitting.** Top: $s^2$ISM-RbN reconstructions of fluorescently labelled F-actin obtained using statistically independent noise realizations generated either experimentally through intra-pixel temporal partitioning of the photon stream (hardware splitting, left) or computationally through stochastic photon splitting of the corresponding photon-counting dataset (software splitting, right). Magnified insets of the same region highlight the close agreement between the two reconstructions in the recovery of fine filamentous structures. Bottom left: pixel-wise difference between the hardware- and software-splitting reconstructions. Bottom right: corresponding absolute normalized difference, obtained after accounting for the overall intensity scale. The asterisk ($^*$) indicates that the displayed maximum corresponds to the 99.9th percentile of the pixel-intensity distribution. Scale bars, 5 $\mu$m.

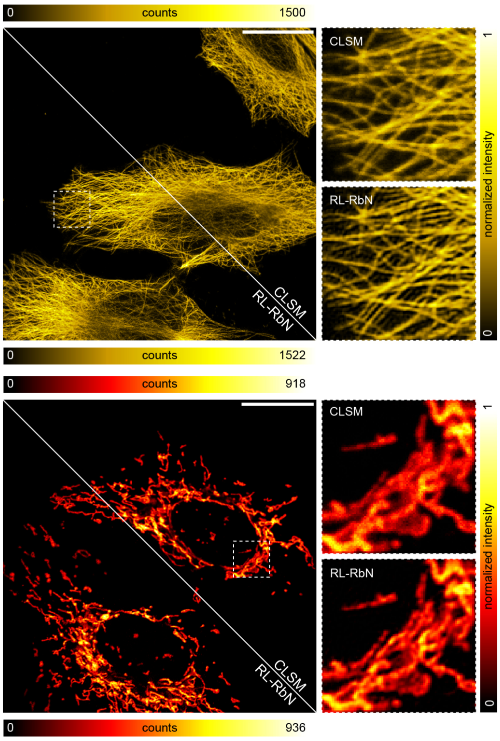


**Figure S14**: **Transfer of RL-RbN to a commercial photon-counting confocal microscope.** Photon-counting CLSM images of $\alpha$-tubulin (top) and TOMM20-labelled mitochondria (bottom), acquired using a commercial Evident microscope, were reconstructed with RL-RbN using computationally generated independent photon-counting realizations. For each target, the full field of view is displayed as a diagonal split between CLSM and RL-RbN, whereas magnified insets highlight the improved separation of densely packed microtubules and the recovery of fine mitochondrial structures without progressive noise amplification. Scale bars: 25 µm for top and 20 µm for bottom.

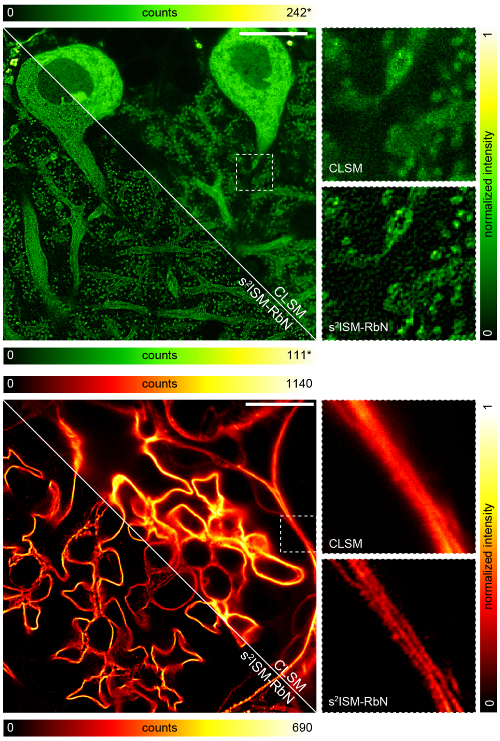


**Figure S15**: **Application of s$^2$ISM-RbN to tissue datasets acquired with a commercial Nikon NSPARC microscope.** Top: EGFP-expressing Purkinje cells in a cleared Thy1-EGFP mouse cerebellum slice. Bottom: mouse kidney cryostat section stained with Alexa Fluor 488-conjugated wheat germ agglutinin, highlighting glomeruli and convoluted tubules. For each specimen, the full field of view is displayed as a diagonal split between the CLSM image and the s$^2$ISM-RbN reconstruction, whereas magnified insets compare the same region and highlight the improved structural definition, optical sectioning and noise suppression achieved by RbN regularization. The asterisk ($*$) indicates that the displayed maximum corresponds to the 99.9th percentile of the pixel-intensity distribution. Scale bars: 15 µm.

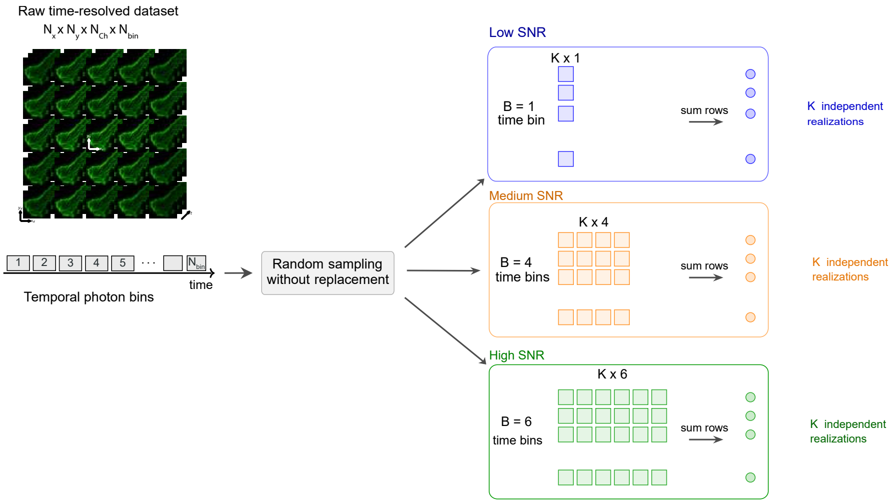


**Figure S16**: **Construction of independent noise realizations from time-resolved SPAD acquisitions.** The raw photon-counting dataset is as a four-dimensional tensor of size $N_x \times N_y \times D \times N_{\text{bin}}$, records, for every scan position and detector channel, the photon counts collected in each of the $N_{\text{bin}}$ temporal bins spanning the pixel dwell time. For each SNR regime, $K \times B$ temporal bins are randomly sampled without replacement and arranged into a $K \times B$ matrix, where $B$ is the number of bins summed to form each realization. Row-wise summation yields $K$ statistically independent noise realizations of the same underlying object. Low-, medium- and high-SNR datasets are generated by using $n_b = 1$, 4, and 6 temporal bins, respectively. Because the subsets of temporal bins are disjoint, the resulting realizations preserve Poisson statistics while remaining statistically independent.
